# Astrocyte glypican 5 regulates synapse maturation and stabilization

**DOI:** 10.1101/2023.03.02.529949

**Authors:** AP Bosworth, M Contreras, S Weiser Novak, L Sancho, IH Salas, U Manor, NJ Allen

**Affiliations:** Molecular Neurobiology Laboratory, The Salk Institute for Biological Studies, 10010 North Torrey Pines Rd, La Jolla, CA, 92037, USA; Neurosciences Graduate Program, University of California, San Diego, La Jolla, CA, 92093, USA; Waitt Advanced Biophotonics Center, The Salk Institute for Biological Studies, 10010 North Torrey Pines Road, La Jolla, CA, 92037, USA

**Keywords:** Astrocyte, synapse, plasticity, development, neurodegeneration

## Abstract

The maturation and stabilization of appropriate synaptic connections is a vital step in the development of neuronal circuits, however the molecular signals underlying these processes are not fully understood. We show that astrocytes, through production of glypican 5 (GPC5), are required for maturation and refinement of synapses in the developing mouse cortex. In the absence of astrocyte GPC5 thalamocortical synapses in the visual cortex show structural immaturity during the critical period, including smaller presynaptic terminals, decreased postsynaptic density area, and presence of more postsynaptic partners at multisynaptic connections. This structural immaturity is accompanied by a delay in developmental incorporation of GLUA2-containing calcium impermeable AMPARs at intracortical synapses. The functional impact of this is that mice lacking astrocyte GPC5 exhibit increased levels of ocular dominance plasticity in adulthood. This shows astrocyte GPC5 is necessary for maturation and stabilization of synaptic connections in typical development, with implications for understanding disorders with altered synaptic function, including Alzheimer’s disease, where GPC5 levels are altered.

## Introduction

The formation and maturation of neuronal synapses during development is essential for circuit function throughout the lifespan. The transformation from immature nascent synapses to stable mature synapses in the adult involves structural changes at the pre and postsynaptic terminals, as well as shifts in neurotransmitter receptor composition. In the cortex functional maturation of excitatory glutamatergic synapses is marked by the incorporation of GLUA2 subunits into AMPA-type glutamate receptors (AMPARs) which renders them impermeable to calcium [1, 2]. Structurally, immature filopodia-like dendritic spines mature into a more stable mushroom-like structure as the synapse is strengthened and stabilized [3]. These processes result in more stable synaptic connections which make up the persistent cortical circuits observed in the adult. The mechanisms which drive this developmental switch in synaptic structure and receptor composition, and that maintain stable connections in the adult brain, are not fully understood.

Astrocytes, a class of glial cell, produce several secreted proteins that regulate the formation and maturation of synapses [4, 5]. Thrombospondins 1 and 2 induce structurally immature excitatory synapses to form, and Hevin regulates postsynaptic spine maturation [6, 7]. Glypicans 4 and 6 induce the formation of nascent glutamatergic synapses by increasing levels of GLUA1 AMPARs, and chordin like 1 (CHRDL1) contributes to synapse maturation by recruitment of GLUA2 AMPARs [8–10]. Glypicans (GPCs) are a family of GPI-anchored heparan sulfate proteoglycans, and there are 6 family members in mammals (GPC1-6) [11]. GPCs exist in a membrane attached form, or are cleaved from the membrane to produce a soluble form, and it is the soluble forms that have been shown to be synaptogenic when produced by astrocytes [8]. GPC family members are expressed in the brain at different stages of development and adulthood by multiple cell types and have roles in regulating synapses [12–14]. Given the synaptogenic role of astrocyte GPC4 and GPC6 in early development, which correlates with their peak of expression [15], we asked if other GPC family members are expressed by astrocytes and if so play a role in regulating synapses. This led us to focus on GPC5, which is expressed widely across the brain, and whose expression by astrocytes is increased during a time of robust synaptic maturation and refinement in the cortex and remains highly expressed in the adult brain [15].

The timing of GPC5 expression suggests it may play a role in the maturation of synapses and their maintenance in the adult, a question we addressed in the mouse visual cortex (VC). Within the primary VC excitatory presynaptic inputs come from two main sources: intracortical and thalamocortical axons. Intracortical inputs can be distinguished by the expression of VGLUT1 at their presynaptic terminals, while thalamocortical inputs, arising from the visual thalamus (dLGN), express VGLUT2 at their presynaptic terminals [16, 17]. Thalamic inputs make up a small fraction of the excitatory connections in the VC but are distinct due to the large size of the axonal boutons and the presence of multiple postsynaptic targets at a single bouton [18–20]. Both intracortical and thalamocortical synapses undergo a developmental incorporation of GLUA2 AMPAR subunits as they mature, which occurs in a layer dependent manner between postnatal (P) day 7-16 [1, 2]. Normal development of the VC involves the refinement of thalamocortical synapses as ocular dominance is established and neurons in the binocular zone (BZ) are tuned for binocular matching of the eyes, which continues during the critical period, a time of enhanced plasticity [21, 22].

Using astrocyte-specific GPC5 conditional knock out (cKO) mice we found that the absence of astrocyte GPC5 renders thalamocortical synapses structurally immature, and delays the incorporation of GLUA2 at intracortical synapses during the critical period, demonstrating that GPC5 regulates synapse maturation. Further, in mice lacking astrocyte GPC5 we found an increase in plasticity in response to visual deprivation in adulthood, showing GPC5 is a plasticity restricting factor in adulthood. The role of GPC5 as a regulator of synapse maturation and plasticity is distinct from the roles of GPC4 and 6 in synapse formation, showing diverse roles for different GPC family members in cortical circuit development and maturation. In humans GPCs have been associated with multiple neurological disorders including autism (GPC4,6), schizophrenia (GPC1,4,5,6), glioma (GPC3), Sanfillipo syndrome type B (GPC5), multiple sclerosis (GPC5) and Alzheimer’s disease (GPC5), demonstrating that understanding the role of GPCs in typical development is important for determining their role in neurological disorders [23–30].

## Results

### *Gpc5* is expressed throughout the brain by both astrocytes and OPCs

We previously used RNA sequencing to analyze the expression of glypican family members by astrocytes in the mouse visual cortex (VC) across postnatal development, with time points correlating with distinct stages of synaptic development: P7 – synapse initiation; P14 – synaptogenesis peaks; P28 – synapse maturation (peak critical period); P120 – synapses stable (adulthood) (Figure 1A) [15]. This showed astrocyte *Gpc5* mRNA is upregulated between P7 and P14, and remains highly expressed throughout adulthood (P28 and P120) (Figure 1B). In contrast, other astrocyte-expressed glypican family members, *Gpc4* and *Gpc6*, peak in expression at P7 and P14 respectively and then decline (Figure 1B). Additionally, the level of *Gpc5* mRNA detected in astrocytes is ∼10-fold higher than either *Gpc4* or *Gpc6* at the peak of expression (Figure 1B), demonstrating *Gpc5* is the predominant glypican expressed by astrocytes. To determine if additional cell types in the mouse cortex express *Gpc5* we consulted published RNA sequencing studies, which show that *Gpc5* mRNA is enriched in both astrocytes and oligodendrocyte progenitor cells (OPCs) compared to other cells including neurons and microglia (Figure 1C) [31]. Thus, *Gpc5* is restricted to the glial lineage.

**Figure 1.**
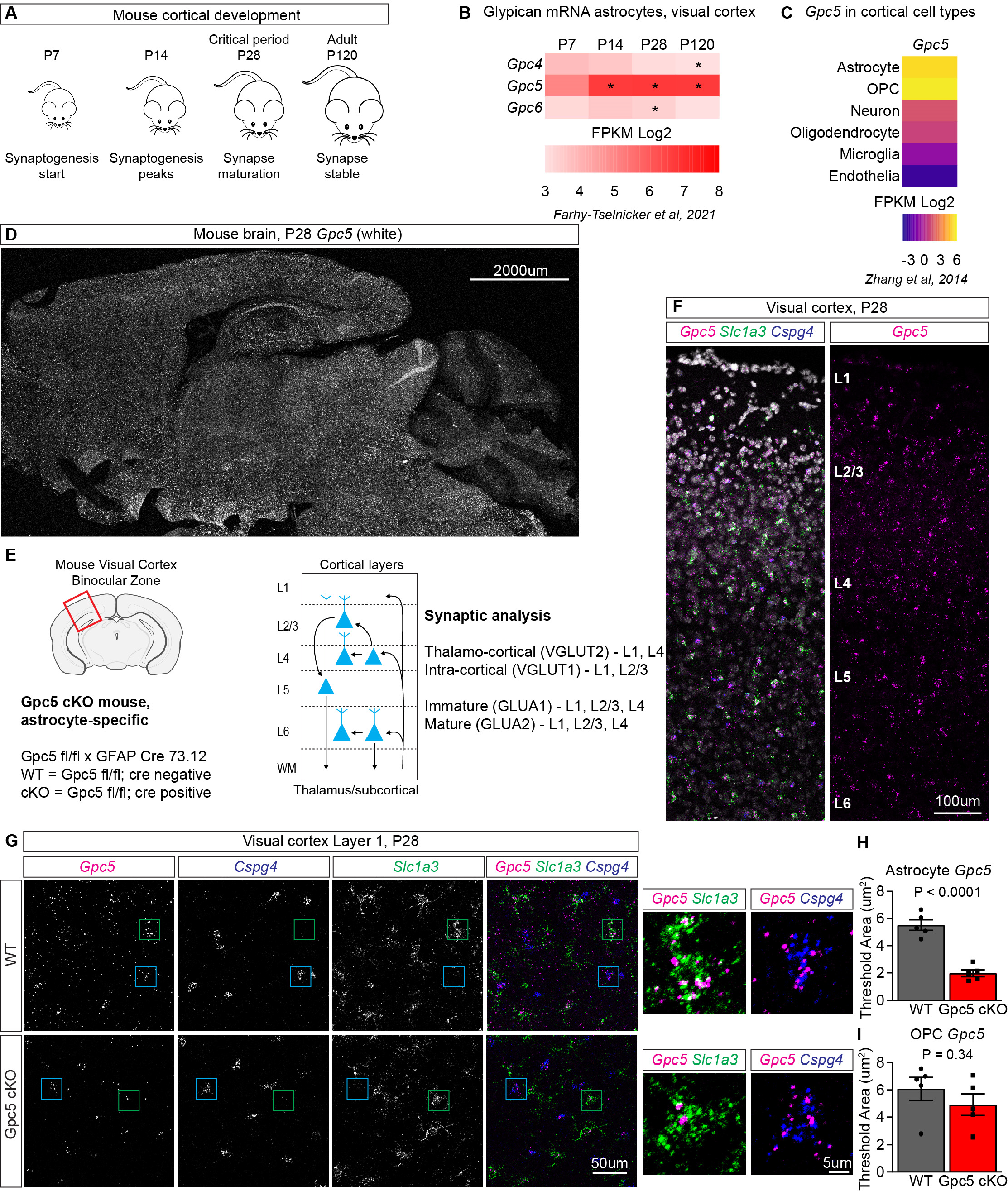
*Gpc5* is expressed throughout the brain by both astrocytes and OPCs. **A.** Mouse cortical synaptogenesis timeline. **B.** Expression of glypican family member mRNA by VC astrocytes across postnatal development shows *Gpc5* increases with age. * denotes significantly altered expression compared to P7. Data from Farhy-Tselnicker et al. 2021. **C.** Cell type expression of *Gpc5* mRNA in the developing mouse cortex shows *Gpc5* is enriched in astrocytes and OPCs. Data from Zhang et al. 2014. **D.** Example image of *Gpc5* mRNA expression in a sagittal section from a WT mouse brain at P28, visualized using smFISH. **E.** Schematic of Gpc5 cKO mouse generation and experiment outline, with synaptic analysis performed in the mouse VC. **F.** Example image of *Gpc5* mRNA expression in both astrocytes (*Slc1a3*) and OPCs (*Cspg4*) in VC at P28 in a WT mouse, visualized using smFISH. **G.** Representative images of WT and Gpc5 cKO P28 L1 VC *Gpc5* mRNA expression, and colocalization with astrocytes (*Slc1a3*) and OPCs (*Cspg4*). Green box shows zoom in image of astrocyte; blue box shows zoom in image of OPC. **H.** *Gpc5* mRNA expression in astrocytes in cKO mice. Quantification of G. **I.** *Gpc5* mRNA expression in OPCs in cKO mice. Quantification of G. H,I: N=5 mice/condition. Graphs show mean ± SEM, individual data points represent mice. Statistics by 2-sided T-test, P-value on graph. See also Figure S1.

To ask how broadly expressed *Gpc5* is in the mouse brain we first consulted published RNA sequencing studies, including our own, that analyzed adult astrocytes from multiple brain regions. These show that *Gpc5* is highly expressed by astrocytes in the forebrain, particularly the cortex, and at a lower level in cerebellar astrocytes (Figure S1A) [32, 33]. As a second approach we performed spatial analysis of *Gpc5* mRNA in sagittal sections of the P28 mouse brain using single molecule fluorescent *in situ* hybridization (smFISH), showing widespread signal across brain regions (Figure 1D). Due to the higher expression of *Gpc5* by cortical astrocytes (Figure S1A), and our time course analysis of *Gpc5* expression in VC astrocytes, we focused on the VC for further studies of *Gpc5*. smFISH of *Gpc5* showed homogeneous expression across all layers of VC at P28, reproducing our published findings (Figure 1F) [15]. Based on the temporal expression of *Gpc5* – upregulated at P14 and remaining high into adulthood, and the known role of other glypican family members in regulating synaptic development, we hypothesized that GPC5 plays a role in regulating synaptic maturation and/or stability.

To ask how astrocyte GPC5 regulates synapses we developed an astrocyte-specific GPC5 knock out mouse by crossing mice with a floxed allele of GPC5 to mice expressing cre recombinase in astrocytes (Gfap-cre 73.12), and compared GPC5_fl/fl_ cre negative (WT) and GPC5_fl/fl_ cre positive (cKO) littermates for all experiments (Figure 1E). To verify astrocyte specific removal of GPC5 we performed smFISH for *Gpc5* in P28 cKO and WT VC along with an astrocyte (*Slc1a3*) and an OPC probe (*Cspg4*) (Figure 1G). In WT mice we detected widespread expression of *Gpc5* throughout the VC in both astrocytes and OPCs as shown by colocalization with the respective cell markers (Figure 1G; S1B). In Gpc5 cKO mice, *Gpc5* expression is significantly decreased in astrocytes but not OPCs demonstrating the specificity of the knockdown (Figure 1H,I). Gpc5 cKO mice retain some *Gpc5* expression in the cortex overall due to OPC expression (Figure S1C). Additionally, we asked whether there is a compensatory response in astrocytes to knocking out GPC5 by probing for two other astrocyte-expressed glypicans, *Gpc4* and *Gpc6,* and found no significant differences (Figure S1D-G). This shows that *Gpc5* is enriched in glial cells in the mouse VC, and that removing GPC5 from astrocytes does not cause a compensatory upregulation of *Gpc5* in OPCs, or *Gpc4* and *Gpc6* in astrocytes.

### Synapses in Gpc5 cKO mice are immature during the critical period

Due to the role of other glypican family members expressed by astrocytes, GPC4 and GPC6, in regulating synaptic development and levels of GLUA1 containing AMPARs, we first asked if GPC5 regulates the number of synapses or their AMPAR composition. Due to the uniform expression of *Gpc5* across upper and lower layers of the VC, and the maintained high expression of *Gpc5* at P28, the peak of the critical period, we analyzed both thalamocortical and intracortical synapses within the VC at this timepoint. To determine the number and AMPAR composition of synapses we used immunohistochemistry to label presynaptic markers VGLUT1 (for intracortical synapses) and VGLUT2 (for thalamocortical synapses), and postsynaptic markers GLUA1 (immature synapses) and GLUA2 (mature synapses), visualized using confocal microscopy (Figure 1E; Figure 2, S2).

**Figure 2.**
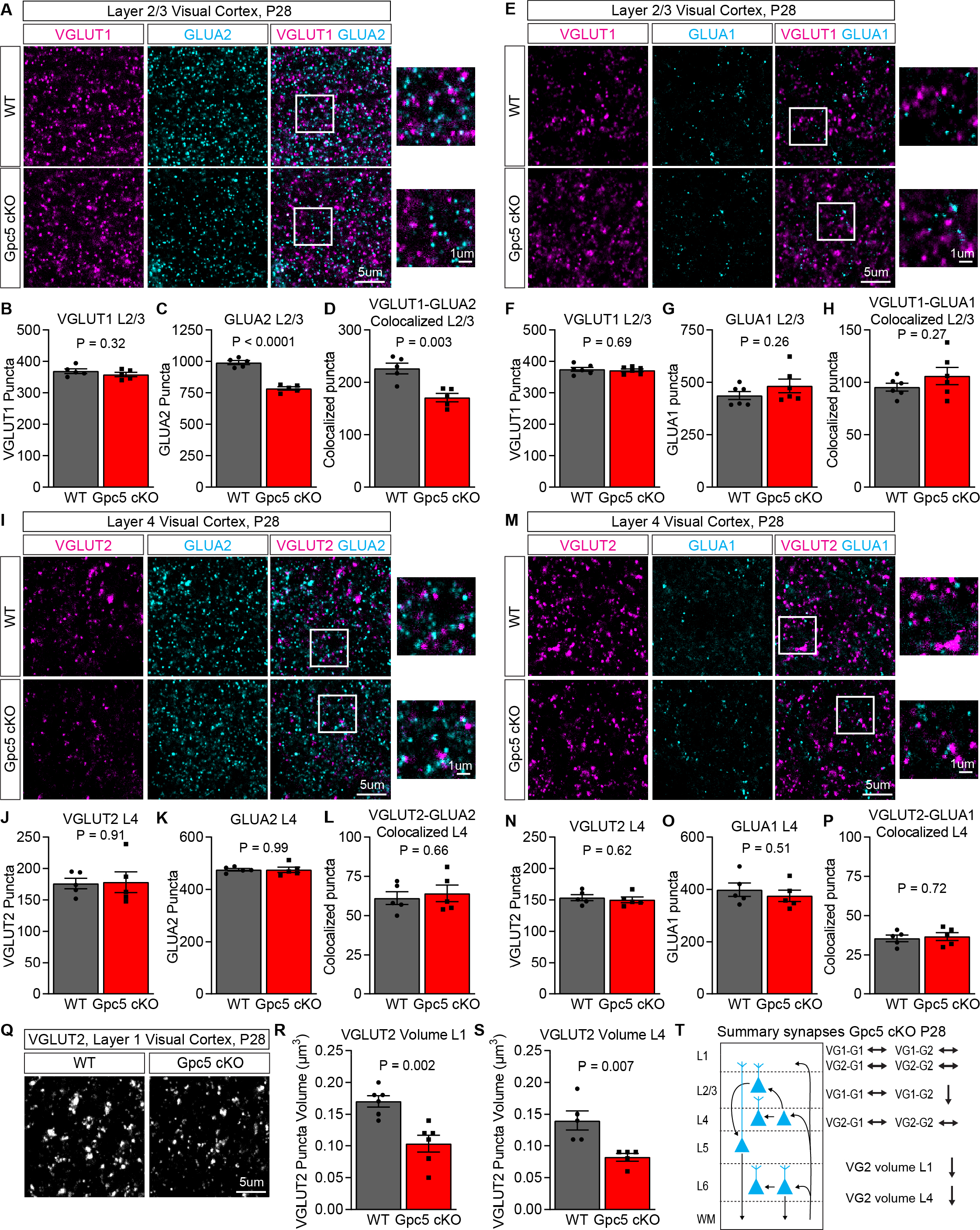
Synapses in Gpc5 cKO mice are immature during the critical period. **A-D.** GLUA2 protein level is decreased at intracortical synapses in Gpc5 cKO mice at P28. **A.** Representative images of immunostaining for intracortical presynaptic marker VGLUT1 and postsynaptic GLUA2 in L2/3. **B-D.** Quantification of immunostaining, number of VGLUT1 (**B**), GLUA2 (**C**) and colocalized (**D**) puncta shows decreased GLUA2 and colocalization. N=5 mice/condition. **E-H.** GLUA1 protein level is unchanged at intracortical synapses in Gpc5 cKO mice at P28. **E.** Representative images of immunostaining for intracortical presynaptic marker VGLUT1 and postsynaptic GLUA1 in L2/3. **F-H.** Quantification of immunostaining, number of VGLUT1 (**F**), GLUA1 (**G**) and colocalized (**H**) puncta shows no change. N=6 mice/condition. **I-P.** Thalamocortical synapses have unaltered AMPAR level in Gpc5 cKO mice at P28. **I.** Representative images of immunostaining for thalamocortical presynaptic marker VGLUT2 and postsynaptic GLUA2 in L4. **J-L.** Quantification of immunostaining, number of VGLUT2 (**J**), GLUA2 (**K**) and colocalized (**L**) puncta shows no change. N=5 mice/condition. **M.** Representative images of immunostaining for thalamocortical presynaptic marker VGLUT2 and postsynaptic GLUA1 in L4. **N-P.** Quantification of immunostaining, number of VGLUT2 (**N**), GLUA1 (**O**) and colocalized (**P**) puncta shows no change. N=5 mice/condition. **Q-S.** VGLUT2 puncta volume is decreased in P28 Gpc5 cKO mice. **Q.** Representative images of VGLUT2 puncta in L1 VC. **R,S.** Quantification of Q, VGLUT2 puncta volume in L1 and L4. N=6 mice/condition L1, N=5 mice/condition L4. **T.** Summary of synaptic changes in Gpc5 cKO mice at P28. Graphs show mean ± SEM, individual data points represent mice. Statistics by 2-sided T-test, P-value on graph. See also Figure S2.

To determine intracortical synapse number and AMPAR composition we analyzed the colocalization of GLUA1 or GLUA2 with the presynaptic marker VGLUT1 in layer 1 (L1) and L2/3 of WT and Gpc5 cKO mice. This showed a significant ∼30% decrease in the colocalization of GLUA2 and VGLUT1 in L2/3, with no difference in L1 (Figure 2A,D; S2A,D). This is driven by a decrease in GLUA2 puncta in L2/3, with no difference in L1, and no alteration in the number of presynaptic terminals marked by VGLUT1 (Figure 2A-C; S2A-C). In the case of GLUA1 we found no difference in the number of GLUA1 puncta in L1 or L2/3, nor in the colocalization of GLUA1 and VGLUT1 in either layer (Figure 2E-H; S2E-H). This shows in the absence of astrocytic GPC5 intracortical synapses lack GLUA2 specifically in L2/3.

To investigate if there are alterations at thalamocortical synapses in Gpc5 cKO mice we quantified the colocalization of the presynaptic marker VGLUT2 with postsynaptic GLUA1 or GLUA2 in L1 and L4, where thalamocortical synapses predominantly form. We found no significant difference in the colocalization between VGLUT2 and GLUA1 in either L1 or L4, or in the number of GLUA1 or VGLUT2 puncta (Figure 2M-P; S2M-P). There is no difference in colocalization of VGLUT2 and GLUA2 in L1 or L4, or in the number of GLUA2 or VGLUT2 puncta, in contrast to the observed decrease in GLUA2 observed at L2/3 intracortical synapses (Figure 2I-L; S2I-L). While the number of presynaptic VGLUT2 terminals is not altered between WT and Gpc5 cKO, we found a significant ∼40% decrease in the volume of VGLUT2 puncta in both L1 and L4 (Figure 2Q-S).

These data show that intracortical and thalamocortical synapses have distinct developmental aberrations in the VC of Gpc5 cKO mice during the critical period (Figure 2T), with decreased GLUA2 at intracortical synapses and apparently smaller presynaptic terminals at thalamocortical synapses. Together this suggests that astrocyte GPC5 contributes to synapse maturation in the developing VC.

### Thalamocortical synapses are structurally immature in Gpc5 cKO mice

The decrease in VGLUT2 puncta volume observed by confocal microscopy in Gpc5 cKO mice suggests a presynaptic structural deficit at thalamocortical synapses. This could be due to a decrease in size of the presynaptic axonal bouton, and/or a decreased recruitment of presynaptic vesicles containing VGLUT2. We investigated this at the ultrastructural level using electron microscopy (EM). To specifically analyze thalamocortical synapses formed between presynaptic VGLUT2 expressing neurons originating in the visual thalamus (dLGN) and L4 target neurons in the VC we used a viral strategy to deliver the EM marker APEX2 to label mitochondria within these axons [34]. We injected AAV9-COX4-DAPEX2 into the dLGN of littermate pairs of WT and Gpc5 cKO mice at P14 and collected brains at P28 for processing for EM, along with treatment with diaminobenzidine (DAB) to visualize APEX2 labeled mitochondria (Figure 3A). Within the VC the APEX2-DAB labeled mitochondria of the dLGN projections were identifiable in L4 where these projections synapse in the VC. Serial sections were collected and imaged in the scanning electron microscope, and high-resolution volumes of neuropil (3DEM) from VC L4 were assembled.

**Figure 3.**
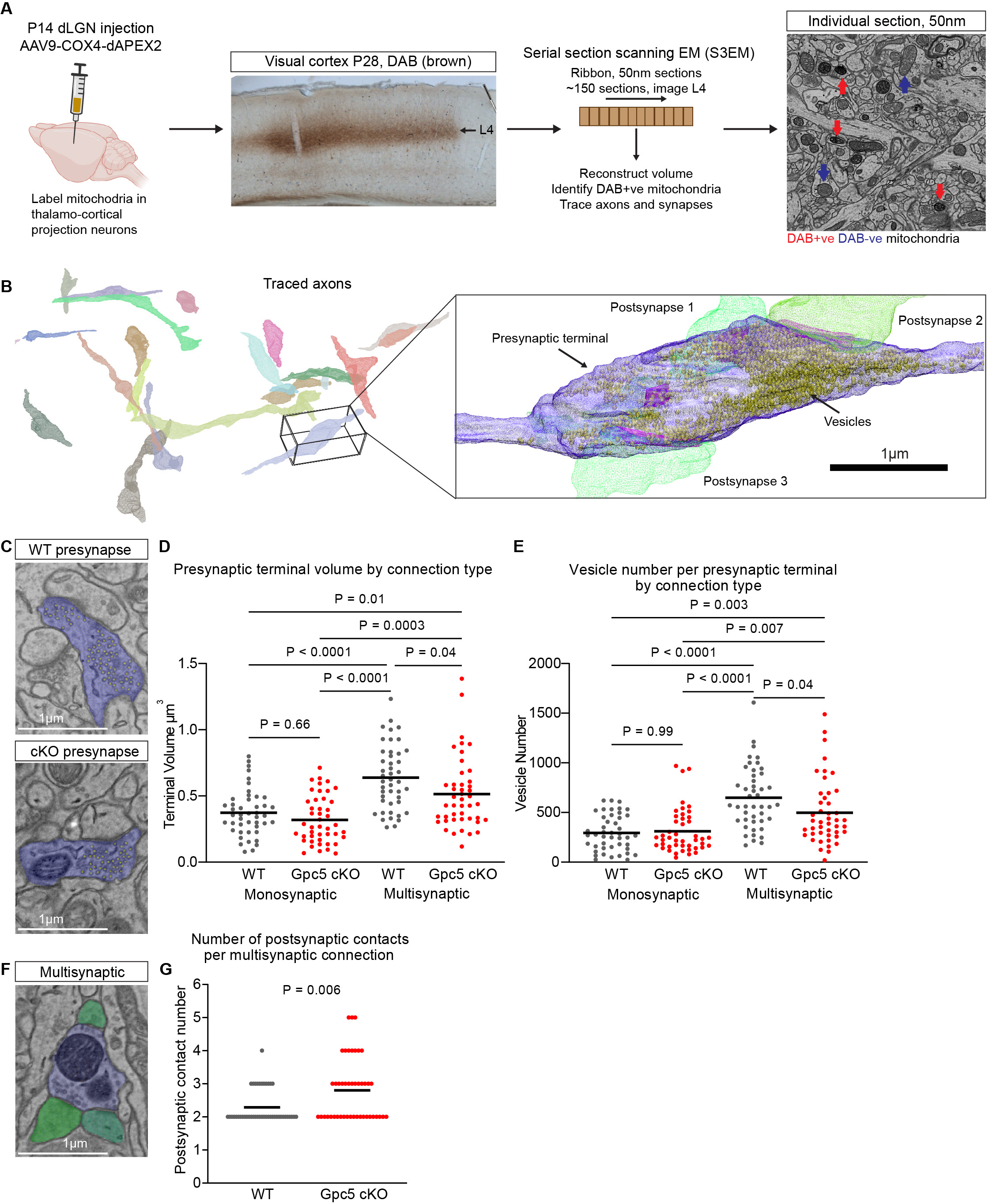
Thalamocortical synapses are structurally immature in Gpc5 cKO mice. **A.** Schematic of experimental design. Mice were injected in the dLGN with AAV9-COX4-DAPEX2 at P14 and collected at P28. Sections of visual cortex underwent a DAB reaction and were processed for EM, with serial sections made at 50nm and loaded in the scanning electron microscope. An ROI in L4 was selected for EM imaging; example single plane image with APEX2+ mitochondria indicated by red arrows, APEX2-mitochondria indicated by blue arrows. **B.** Example of reconstructed APEX2 positive thalamocortical axons in L4. Zoom in: Reconstructed thalamocortical presynaptic bouton and postsynaptic spines. Bouton in purple, vesicles in yellow, spines in green, PSD in magenta. **C.** Example images of WT and Gpc5 cKO thalamocortical presynaptic boutons. Vesicles are marked in yellow, bouton shaded purple. **D.** Volume of multisynaptic, but not monosynaptic, thalamocortical axonal boutons are decreased in Gpc5 cKO mice compared to WT. **E.** The number of synaptic vesicles in multisynaptic thalamocortical axonal boutons is decreased in Gpc5 cKO mice compared to WT. D,E graphs show mean, individual data points represent presynaptic boutons. N=45 presynaptic boutons per condition, statistics by two-way ANOVA, P-values on graph. **F.** Example image of multisynaptic connection, presynaptic bouton shaded purple and postsynaptic spines green. **G.** The number of postsynaptic contacts at multisynaptic thalamocortical boutons is increased in Gpc5 cKO mice. Graph shows mean; individual data points represent presynaptic boutons. N=45 presynaptic boutons per condition, statistics by Mann-Whitney test, P-value on graph. See also Figure S3.

Thalamocortical presynaptic boutons were identified by the presence of DAB labeled mitochondria within the parent axon, and thalamic connections traced and reconstructed (Figure 3B). Labeled boutons had the features of VGLUT2 positive thalamic synapses, including a large volume, asymmetric synapses, and multiple postsynaptic contacts at some presynaptic sites. We analyzed a number of features at each reconstructed presynaptic bouton, including bouton volume, number of synaptic vesicles and number of postsynaptic partners. In each analysis we compared features of WT and Gpc5 cKO synapses as a single group, as well as analyzing monosynaptic and multisynaptic connections as separate groups.

We found the average volume of presynaptic boutons in Gpc5 cKO mice is significantly decreased compared to WT (Figure S3A). This result is driven by a significant decrease in the volume of multisynaptic boutons in the Gpc5 cKO, with no difference in the volume of monosynaptic boutons (Figure 3C,D). In both genotypes there is a significant increase in the volume of multisynaptic boutons compared to monosynaptic, as is expected, but this increase is smaller in the Gpc5 cKO (Figure 3D). The number of synaptic vesicles within multisynaptic boutons is significantly decreased in the Gpc5 cKO compared to WT, with no change at monosynaptic connections (Figure 3C,E; S3B). Additionally, we found that the average number of postsynaptic partners at a multisynaptic bouton is higher in the Gpc5 cKO compared to WT, and that the maximum observed number of synapses at a single bouton is also greater in the Gpc5 cKO (Figure 3F,G).

These results demonstrate that there are structural alterations at thalamocortical synapses in Gpc5 cKO mice which are particularly pronounced at multisynaptic connections. The decreased VGLUT2 puncta volume observed by light microscopy (Figure 2Q-S) is likely a result of the decreased volume of Gpc5 cKO multisynaptic thalamic boutons and decreased total number of vesicles per bouton. These alterations, as well as the larger number of postsynaptic contacts at multisynaptic boutons, suggests that the absence of astrocytic GPC5 renders thalamocortical synapses more immature [3, 35, 36].

### Gpc5 cKO mice show altered postsynaptic structure at thalamocortical synapses

Our data demonstrate that there are developmental disruptions in the presynaptic structure of thalamocortical synapses and the synaptic AMPAR composition of intracortical synapses in the absence of GPC5 (Figure 2,3). We next asked whether these effects are accompanied by a change in postsynaptic structure of either thalamocortical or intracortical synapses.

To specifically analyze postsynaptic ultrastructure of thalamocortical synapses we used the EM dataset described above and segmented the postsynaptic partners of the APEX2 labeled presynaptic boutons that had been reconstructed. The predominant postsynaptic structures identified were dendritic spines, with labeled synapses directly onto the dendritic shaft rarely observed. To characterize the postsynaptic compartment, we first measured the surface area of the postsynaptic density (PSD), finding a significant decrease in PSD surface area in Gpc5 cKO mice (Figure 4A,B; S4A). This decrease is present at spines opposed to both monosynaptic and multisynaptic boutons (Figure 4B). Given the decreased size of the PSD we asked if postsynaptic spine structure is shifted to a more immature phenotype. We classified reconstructed spines as thin, mushroom or other based on morphology (see Methods). When analyzing all connections together we found a trend towards an increased percentage of thin spines and a decreased percentage of mushroom spines in the Gpc5 cKO (Figure S4B), which became significant when spines were separated into those present at monosynaptic or multisynaptic connections (Figure S4C). To ask which connection type is responsible for this difference we separately analyzed spines opposing monosynaptic and multisynaptic boutons. This showed that there is a significant increase in the percentage of thin spines and decrease in mushroom spines at monosynaptic connections (Figure 4C), with no significant shift at multisynaptic connections in Gpc5 cKO mice (Figure 4D).

**Figure 4.**
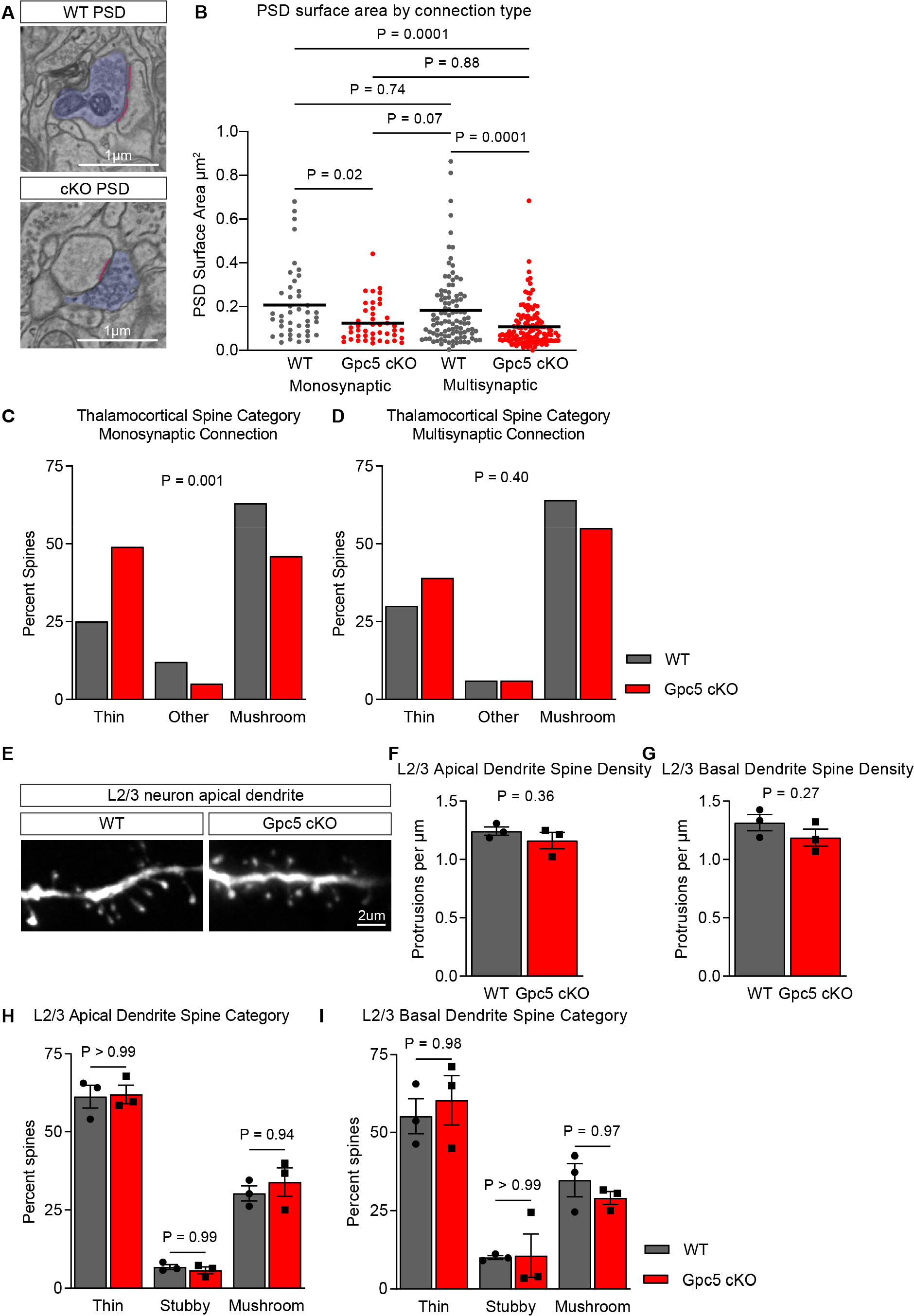
Gpc5 cKO mice show altered postsynaptic structure at thalamocortical synapses. **A-D.** Postsynaptic structures at L4 thalamocortical synapses are immature in Gpc5 cKO mice. **A.** Example images of WT and Gpc5 cKO PSDs in L4. PSD marked in magenta, axon in purple. **B.** Surface area of PSD at APEX2 positive thalamocortical synapses is decreased at both monosynaptic and multisynaptic connections in Gpc5 cKO mice. Graphs show mean, individual data points represent PSDs. N=45 presynaptic boutons and associated PSDs per condition, statistics by two-way ANOVA, P-values on graph. **C,D.** Dendritic spine morphologies are shifted towards a more immature state in Gpc5 cKO mice. Prevalence of thin spines is increased and mushroom spines decreased at monosynaptic (**C**) but not multisynaptic (**D**) APEX2 positive synapses. Other category includes stubby spines and synapses directly onto the dendritic shaft. Data presented as percentage of spines in each category, spines were identified as compartments opposed to presynaptic boutons analyzed in Figure 3. Statistics by Chi-square test, P-value on graph. **E-I.** There are no gross morphological changes to dendritic spines of P28 L2/3 pyramidal cells in Gpc5 cKO mice. **E.** Representative images of P28 L2/3 pyramidal cell dendrites. **F,G.** Quantification of spine density for apical (**F**) and basal (**G**) dendrites respectively. **H,I.** Categorization of spine shape for secondary apical (**H**) and basal (**I**) dendrites respectively. N=3 mice. Graphs show mean ± SEM, individual data points represent mice. G,H statistics by T-test, P-value on graph; I,J statistics by two-way ANOVA, P-value on graph. See also Figure S4.

Given the decreased GLUA2 levels present at L2/3 intracortical synapses in the P28 Gpc5 cKO mice, we next asked if there was a shift towards a more immature dendritic spine structure in neurons in this layer, which consists of predominantly intracortical synapses. To investigate this, we used sharp electrodes to fill individual L2/3 pyramidal neurons in the VC with fluorescent dye in acute brain sections of P28 WT and Gpc5 cKO mice, with spines imaged using confocal microscopy (Figure 4E). We quantified spine density, spine length, spine head diameter, and head/neck ratio of spines located on secondary apical and basal dendrites. We found no significant difference in the average spine density on apical or basal dendrites (Figure 4F,G). Analysis of spine morphology (width, length, length to width ratio) also showed no difference between genotypes (Figure S4D-I). We further categorized spines as mushroom, thin or stubby based on these measurements, finding no differences in categorization between the genotypes for either apical or basal dendrites (Figure 4H,I). Overall, we did not observe any gross changes in the structure of spines on dendrites of L2/3 neurons in Gpc5 cKO mice.

The decreased PSD surface area at thalamocortical synapses in Gpc5 cKO mice indicates that the strength of thalamocortical synapses is diminished, while the shift towards an increased prevalence of immature thin spines suggests that thalamocortical synapses are more immature. This immature postsynaptic phenotype, along with the smaller and less refined thalamic axonal boutons described above, indicate that the absence of astrocytic GPC5 significantly disrupted the maturation of thalamocortical synapses.

### GPC5 is sufficient to induce presynaptic specializations

Analysis of synapses in the VC of Gpc5 cKO mice showed a number of alterations, including structurally immature thalamocortical synapses and decreased GLUA2 at intracortical synapses. To gain insight into the site of action of GPC5, i.e. presynaptic or postsynaptic, we performed experiments using retinal ganglion cell (RGC) neurons in culture to ask if soluble GPC5 is sufficient to induce synapses to form. We used RGCs as they form few synapses in the absence of astrocytes, and have successfully been used to study the role of astrocytes and astrocyte-secreted proteins in synaptogenesis, including GPC4 and GPC6 [4].

RGCs were cultured alone, with astrocytes or with recombinant GPC5 protein added to the media for 6 days, then immunostained for presynaptic VGLUT2 and postsynaptic PSD95 (Figure 5A). Analysis of colocalized VGLUT2 and PSD95, to mark synapses, showed a significant increase in synapse number in RGCs cultured with astrocytes compared to alone, with no effect of soluble GPC5 on synapse number (Figure 5B). This demonstrates that GPC5 is not sufficient to induce synapse formation by itself. Due to the effect of GPC5 on presynaptic terminals identified in Gpc5 cKO mice, we analyzed whether soluble GPC5 protein was sufficient to increase the number or size of VGLUT2 presynaptic specializations. This identified that GPC5 significantly increased the number of presynaptic sites compared to RGCs cultured alone, though this increase was less than that induced by astrocytes (Figure 5C). Analysis of the size of presynaptic terminals found no significant increase after treatment with GPC5, whereas astrocytes did induce larger terminals (Figure 5D; S5A). This suggests that the site of action of soluble GPC5 is presynaptic, the same as for other astrocyte-expressed glypicans including GPC4 [9].

**Figure 5.**
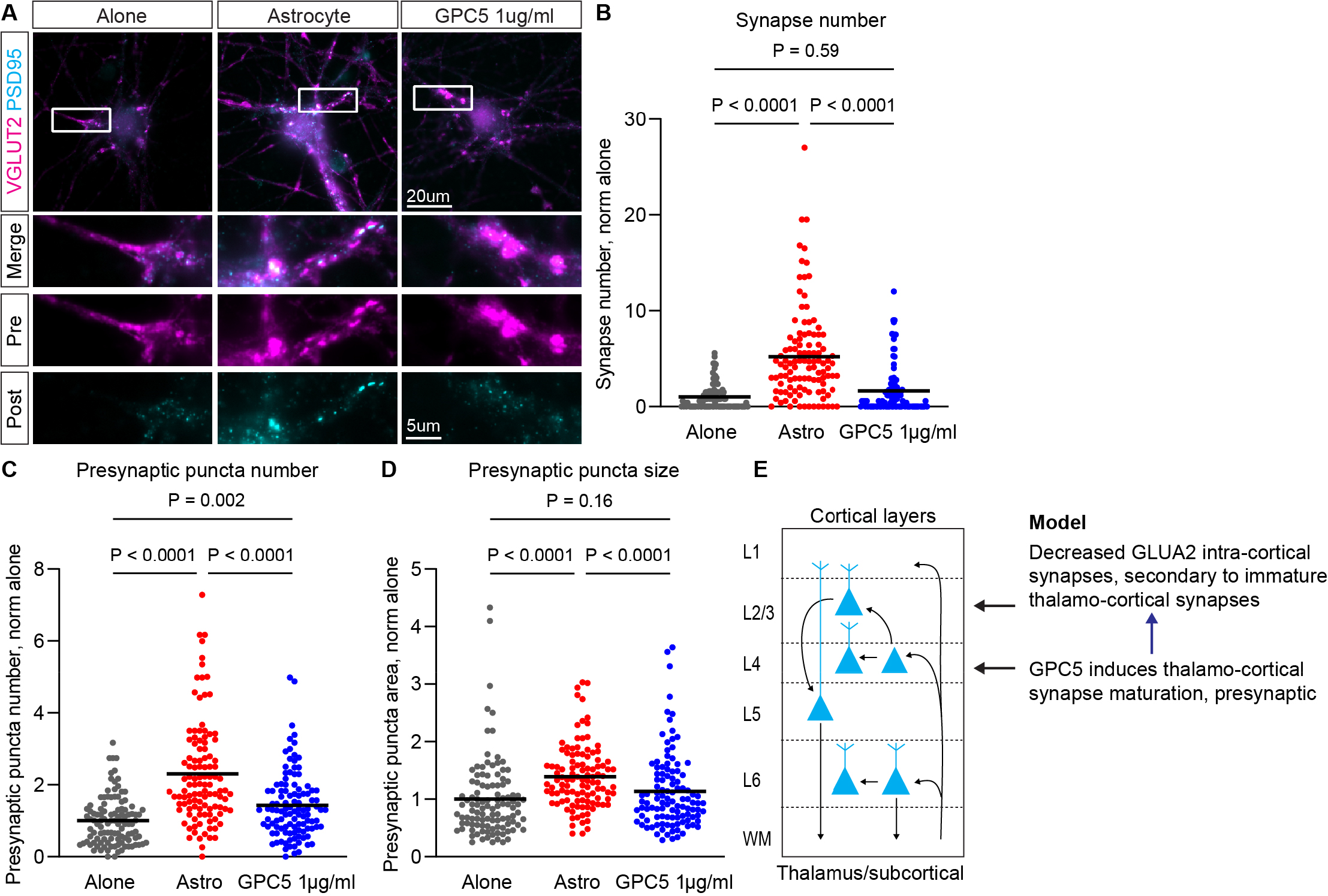
GPC5 is sufficient to induce presynaptic specializations. **A-D.** Treating RGC neurons in culture is sufficient to increase the number of presynaptic terminals without increasing synapse number. **A.** Example images of neurons immunostained for VGLUT2 and PSD95, grown alone, with astrocytes or GPC5 protein. **B.** Quantification of synapse number. **C.** Quantification of presynaptic terminal number. **D.** Quantification of presynaptic terminal size. Graphs show mean, individual data points represent neurons. N=110 cells per condition from 4 experiments, statistics by Kruskal-Wallis ANOVA on ranks with Dunn’s post-hoc test, P-values on graph. **E.** Model of GPC5 action. See also Figure S5.

Based on these findings we hypothesize that *in vivo* the primary target of astrocyte GPC5 is presynaptic thalamocortical axonal boutons that synapse onto L4 neurons. As L4 neurons project to L2/3 neurons, we hypothesize that immature thalamocortical synapses have downstream effects that impact GLUA2 AMPAR levels at intracortical synapses (Figure 5E).

### Large scale ocular dominance plasticity during the critical period is unchanged in Gpc5 cKO mice

Gpc5 cKO mice show features of immature synapses at P28, the peak of the critical period, namely immature synapse structure at thalamocortical synapses and decreased GLUA2 at intracortical synapses. This led us to ask whether Gpc5 cKO mice show enhanced experience dependent plasticity in the VC during the critical period, a time when plasticity is already high and brief periods of sensory deprivation are sufficient to alter neuronal connectivity. In the binocular zone (BZ) of the VC this plasticity can be observed by depriving one eye of vision for a number of days, which induces neurons from the open eye to expand their territory in the BZ.

To assess this Gpc5 cKO and WT mice were monocularly enucleated (ME) at P28 and the extent of BZ remodeling assessed after 12 hours (baseline innervation) or 5 days (remodeling) by exposing mice to bright light to activate neurons in the VC and induce expression of the immediate early gene *Arc,* visualized using smFISH (Figure 6A) [37]. The width of the *Arc* signal represents the BZ innervated by the intact eye, and expansion of the *Arc* signal following ME provides a measurement of ocular dominance plasticity. After 12 hours of deprivation, which represents baseline innervation of the BZ by the non-deprived eye, we found no difference in the width of the *Arc* activated neuron zone between Gpc5 cKO mice and WT indicating that absence of astrocyte GPC5 does not alter baseline connectivity (Figure 6B,C). In both the WT and Gpc5 cKO mice, 5 days of ME is sufficient to significantly increase the width of the *Arc* signal compared to 12 hours, indicating remodeling has occurred. Furthermore, we found no significant difference between WT and Gpc5 cKO mice in the width of the *Arc* signal following 5 days of deprivation (Figure 6B,C). This demonstrates that lack of GPC5 in astrocytes does not affect large scale sensory remodeling during the critical period.

**Figure 6.**
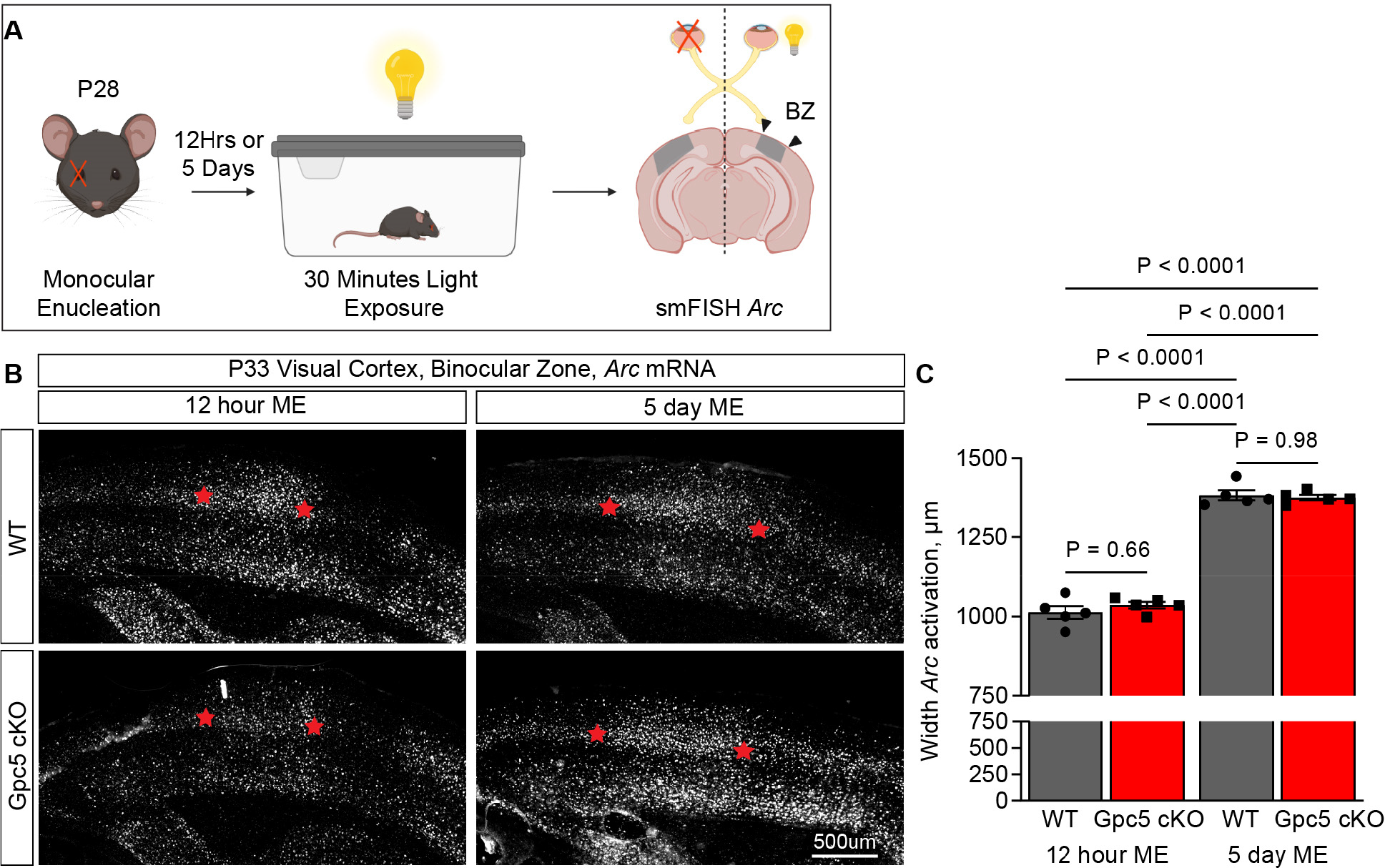
Large scale ocular dominance plasticity during the critical period is unchanged in Gpc5 cKO mice. **A.** Schematic of experiment. P28 WT and Gpc5 cKO mice underwent monocular enucleation (ME) and were collected after 12 hours or 5 days following exposure to bright light. Sections of VC were probed for *Arc* mRNA using smFISH to visualize activated neurons. **B.** Representative images of *Arc* mRNA in VC ipsilateral to nondeprived eye in WT and Gpc5 cKO mice, 12 hours or 5 days after ME. **C.** Ocular dominance plasticity is unchanged in P28 Gpc5 cKO mice compared to WT, quantification of B. Graph mean ± SEM, individual data points mice. N=5 mice/condition, statistics by two-way ANOVA, P-values on graph.

### Synapse maturation is delayed in Gpc5 cKO mice

As *Gpc5* remains highly expressed in the adult brain we asked if adult mice lacking GPC5 maintain deficits in synaptic AMPAR composition and presynaptic terminal size that are present during the critical period (Figure 2). We assessed this using immunohistochemistry and confocal imaging of synapses in the VC of Gpc5 cKO and WT mice at P120. We analyzed intracortical and thalamocortical synapses using the presynaptic markers VGLUT1 and VGLUT2 respectively and focused on the AMPAR subunit GLUA2 due to the decreased level we observed at P28.

At intracortical synapses we found no significant difference in VGLUT1 or GLUA2 puncta number, or colocalization of VGLUT1 and GLUA2, between WT and Gpc5 cKO mice in either L1 or L2/3, although there is a non-significant trend to decreased VGLUT1-GLUA2 in L1 (Figure 7A-H). This is in contrast to P28 where total GLUA2 and VGLUT1-GLUA2 synapses are decreased in L2/3 in the Gpc5 cKO (Figure 2). At thalamocortical synapses we found no difference in VGLUT2 or GLUA2 puncta number, or colocalization of VGLUT2 and GLUA2, between WT and Gpc5 cKO mice in either L1 or L4, although there is a non-significant trend to decreased VGLUT2 puncta number in L4 (Figure 7I-P). This is consistent with findings at P28 (Figure 2). We found no difference in the volume of VGLUT2 puncta between WT and Gpc5 cKO mice in either L1 or L4 (Figure 7Q-S), in contrast to P28 where terminal volume is decreased in Gpc5 cKO (Figure 2).

**Figure 7.**
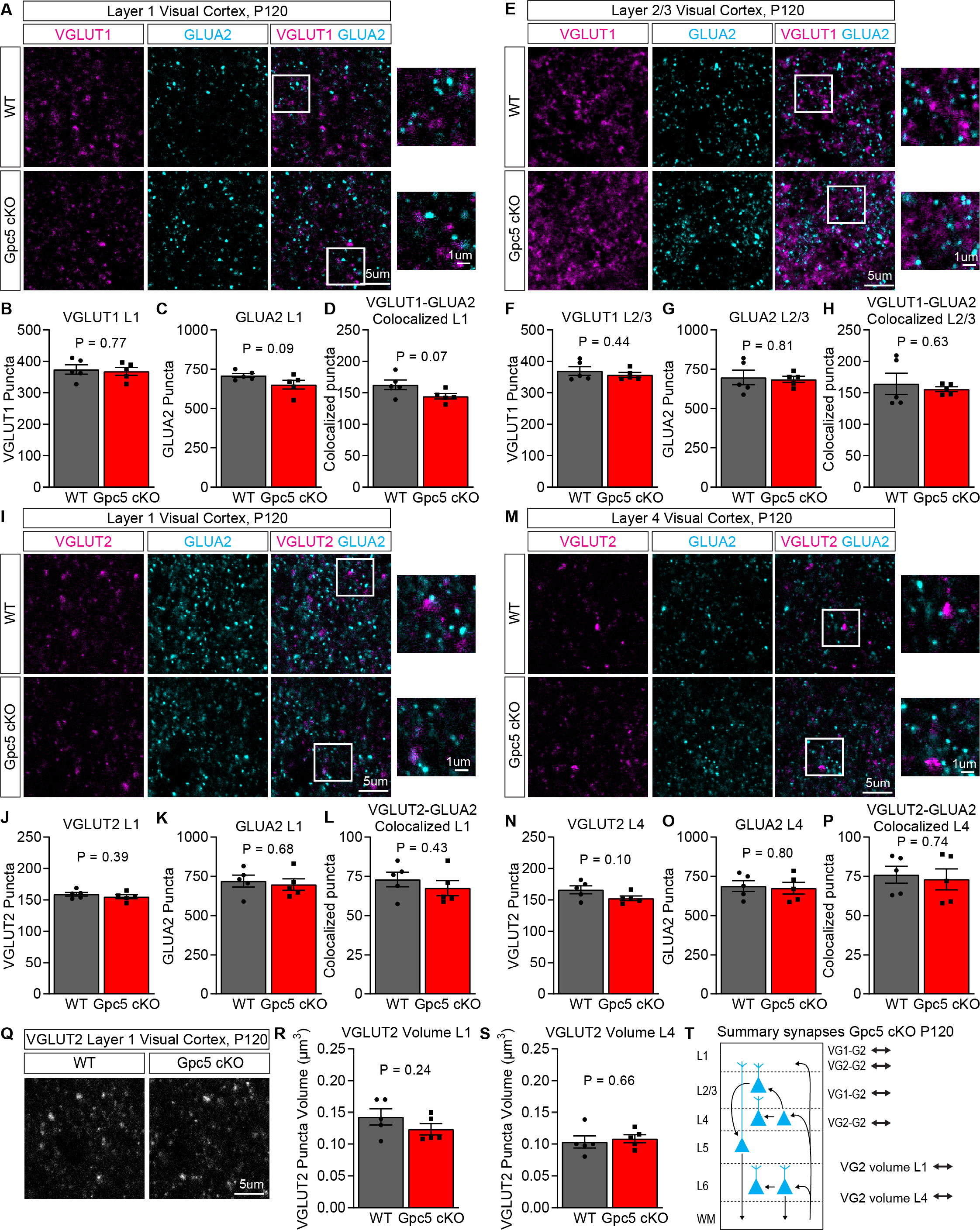
Synapse maturation is delayed in Gpc5 cKO mice. **A-H.** GLUA2 puncta number is recovered at intracortical synapses in Gpc5 cKO mice at P120. **A,E.** Representative images of immunostaining for intracortical presynaptic marker VGLUT1 and postsynaptic GLUA2 in L1 (**A**) and L2/3 (**E**). **B,F.** VGLUT1 puncta number is unchanged in L1 and L2/3. **C,G.** GLUA2 puncta number is unchanged in L1 and L2/3. **D,H.** Colocalization of VGLUT1 and GLUA2 puncta is unchanged in L1 and L2/3. **I-P.** Thalamocortical synaptic levels of GLUA2 are unaltered at P120 in Gpc5 cKO mice. **I,M.** Representative images of immunostaining for thalamocortical presynaptic marker VGLUT2 and postsynaptic GLUA2 in L1 (**I**) and L4 (**M**). **J,N.** VGLUT2 puncta number is unchanged in L1 and L4. **K,O.** GLUA2 puncta number is unchanged in L1 and L4. **L,P.** Colocalization of VGLUT2 and GLUA2 is unchanged in L1 and L4. **Q-S.** VGLUT2 puncta volume is recovered at P120 in Gpc5 cKO mice. **Q.** Representative images of VGLUT2 puncta in layer 1 VC. **R,S.** Quantification of Q, VGLUT2 puncta volume in L1 (**R**) and L4 (**S**). All experiments: N=5 mice/condition. Graphs show mean ± SEM, individual data points mice. Statistics by T-test, P-values on graph. **T.** Summary of synaptic changes in Gpc5 cKO mice at P120.

Taken together this data indicates that by P120 most synaptic alterations detected during the critical period in Gpc5 cKO mice have been rectified (Figure 7T). This suggests that absence of astrocytic GPC5 delays rather than prevents synapse maturation.

### Absence of astrocyte GPC5 enables enhanced ocular dominance plasticity in adulthood

In adulthood the high level of experience dependent plasticity present during the critical period is decreased, and brief periods of sensory deprivation are insufficient to induce large scale remodeling. Although the steady state synapse number and AMPAR composition in Gpc5 cKO mice has mostly reached WT levels in adulthood (Figure 7), the continued high expression of GPC5 by astrocytes in the adult brain (Figure 1B) led us to ask if absence of GPC5 enables increased ocular dominance plasticity in Gpc5 cKO mice in adulthood.

To assess this, we performed monocular enucleation (ME) in Gpc5 cKO and WT mice at P120, and probed for changes in VC neural connectivity using smFISH for *Arc* to visualize active neurons, as described at P28 (Figure 8A). We found no difference in the width of the *Arc* zone between WT and Gpc5 cKO mice after 12 hours of ME indicating no baseline changes in connectivity are present at P120 (Figure 8B,C). After 5 days of ME, we found no difference in the width of the *Arc* zone in WT mice when compared to 12 hours deprivation, consistent with limited plasticity present in the adult brain (Figure 8B,C). For Gpc5 cKO mice we found a significant increase in the width of the *Arc* zone after 5 days of ME compared to 12 hours, demonstrating that absence of astrocytic GPC5 has enabled some plasticity to occur (Figure 8B,C). This was also reflected in the width of the *Arc* zone after 5 days ME being significantly larger in the Gpc5 cKO compared to WT (Figure 8B,C).

**Figure 8.**
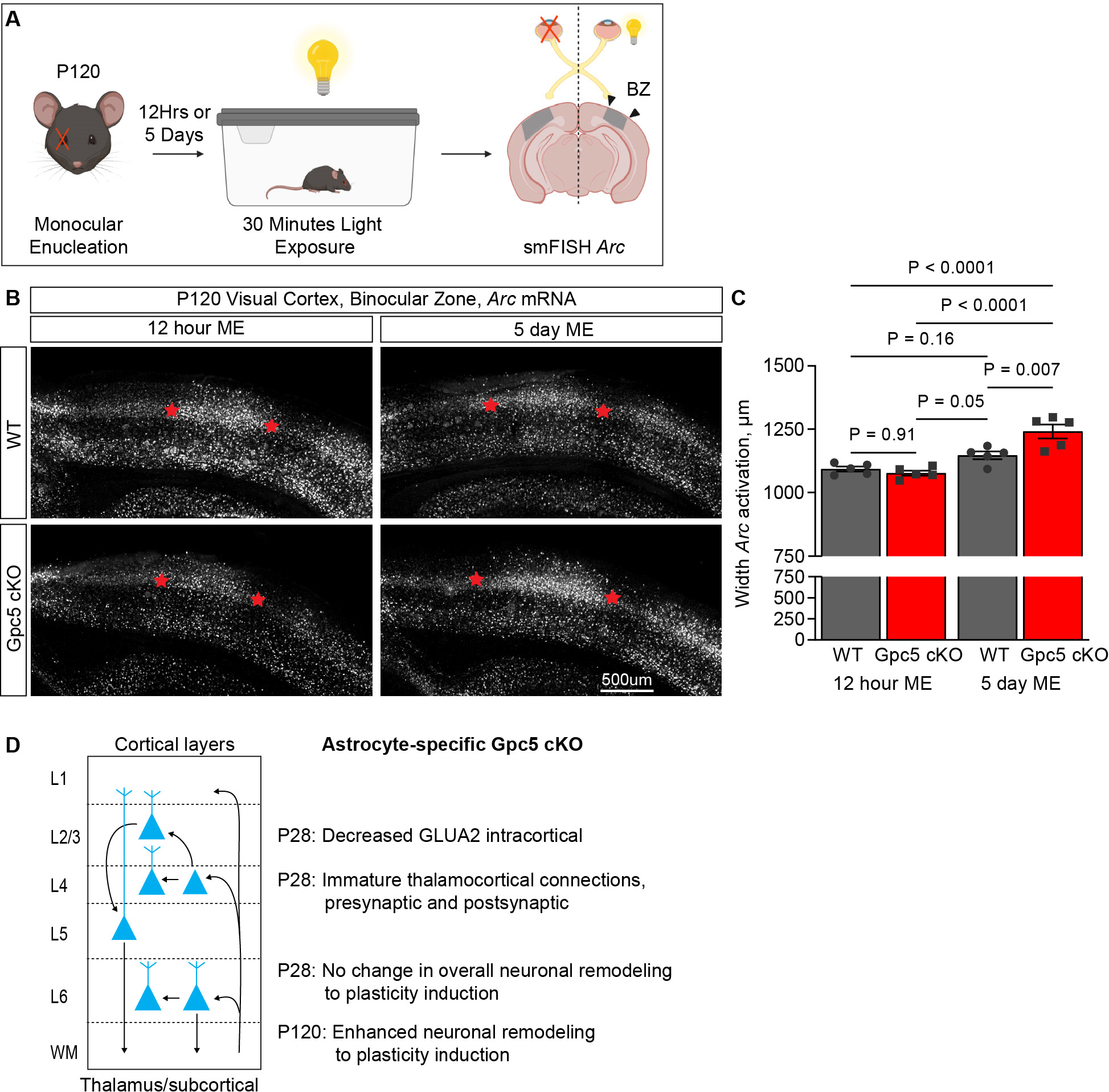
Absence of astrocyte GPC5 enables enhanced ocular dominance plasticity in adulthood. **A.** Schematic of experiment. P120 WT and Gpc5 cKO mice underwent monocular enucleation (ME) and were collected after 12 hours or 5 days following exposure to bright light. Sections of VC were probed for *Arc* mRNA using smFISH to visualize activated neurons. **B.** Representative images of *Arc* mRNA in VC ipsilateral to nondeprived eye in WT and Gpc5 cKO mice, 12 hours or 5 days after ME. **C.** Ocular dominance plasticity is enhanced in P120 Gpc5 cKO mice compared to WT. Quantification of B. Graph mean ± SEM, individual data points mice. N=5 mice/condition, statistics by two-way ANOVA, P-values on graph. **D.** Summary of phenotypes in GPC5 astrocyte-specific cKO mice.

This demonstrates that lifelong absence of GPC5 specifically in astrocytes enables an environment that is permissive to plasticity and neuronal remodeling in the adult brain (Figure 8D).

## Discussion

In this study we identified that astrocyte GPC5 regulates excitatory synapse maturation and stabilization. In the absence of GPC5 in astrocytes the structural maturation and refinement of thalamocortical synapses is impaired, and the level of GLUA2 AMPARs at intracortical synapses is reduced. The consequence of this is enhanced remodeling of connections in the VC after visual deprivation in the adult brain, but not during the critical period, suggesting that in the adult brain GPC5 represses plasticity. Importantly these effects are observed in the presence of unaltered GPC5 expression in OPCs, suggesting there may be distinct roles for GPC5 depending on the cell-type of origin. These actions are also distinct from those of other astrocyte-expressed GPC family members. GPC4 and GPC6 induce nascent synapse formation via clustering of GLUA1 AMPARs, distinct from GPC5 which regulates synapse maturation [8, 9]. Other astrocytic factors that regulate synapse maturation also have distinct actions, for example CHRDL1 recruits GLUA2 AMPARs to thalamocortical synapses, and Hevin regulates NMDA receptors and spine maturation [6, 10]. This shows that while astrocytes produce multiple synapse-regulating factors, including GPC5, each has a distinct action in regulating the development and maturation of excitatory synapses.

We found that astrocyte GPC5 regulates thalamocortical synapse maturation, with these connections showing multiple indications of weaker synapses including smaller bouton size, fewer synaptic vesicles and smaller PSD surface area in cKO mice [36, 38, 39]. The presence of weaker synapses in conjunction with a larger number of postsynaptic partners at multisynaptic boutons suggests that this circuit is not undergoing typical maturation, whereby selected synapses are strengthened and stabilized and excess synapses are pruned [3, 18]. Based on our findings we hypothesize that GPC5 is necessary for the strengthening of appropriate synapses. In the absence of this strengthening, pruning may be disrupted leading to excessive weaker synaptic connectivity particularly at multisynaptic boutons. Weaker synapses with more immature dendritic spine morphologies are indicative of a less stable synaptic connection, and in the adult destabilized axonal boutons have been associated with cognitive decline [40]. Diminished GPC5 expression may therefore contribute to synapse loss via destabilization of axonal boutons, which is of interest as decreased GPC5 expression has been associated with Alzheimer’s disease, where synapse loss is an early pathology [24].

In the absence of GPC5 in astrocytes brief visual deprivation in adulthood is sufficient to induce plasticity, whereas absence of GPC5 does not enhance the level of plasticity that normally occurs during the critical period. This suggests that GPC5 may be involved in actively repressing plasticity after the closure of the critical period. Astrocyte secreted factors, such as CHRDL1, have been shown to repress plasticity through the recruitment of GLUA2 AMPAR subunits [10]. Although Gpc5 cKO mice have reduced synaptic GLUA2 during the critical period, they appear to have recovered GLUA2 levels in the adult, so it is unlikely that this is the mechanism through which GPC5 represses plasticity. The neuronal factor PIRB also represses plasticity, and increased spine density has been found in these KO mice [41]. This is of interest due to increased spine number observed at multisynaptic thalamocortical boutons in Gpc5 cKO mice during the critical period. In the future it will be important to determine if GPC5 is actively repressing plasticity through direct regulation of synaptic stability or AMPAR subunit composition. Alternatively, increased plasticity in adulthood may be the result of incomplete circuit maturation and represent incomplete closure of the critical period in Gpc5 cKO mice.

Identifying the mechanism of how astrocytic GPC5 regulates thalamocortical synapses will help elucidate whether the lack of thalamocortical refinement in the cKO is the result of diminished synaptic pruning, aberrant synapse formation, or failure of synaptic strengthening and stabilization. Our cell culture experiments demonstrated that soluble GPC5 protein is sufficient to induce the formation of presynaptic specializations, without increasing synapse number, suggesting that the primary action of GPC5 is to regulate presynaptic maturation. Based on this we hypothesize that the observed decrease in GLUA2 at intracortical synapses in L2/3 is secondary to a failure of thalamocortical synapses in L4 to mature, as many L4 neurons synapse directly onto L2/3 neurons. *Gpc5* is homogenously expressed by astrocytes across all cortical layers, and yet the major structural phenotypes we identified in Gpc5 cKO mice were at thalamocortical connections. This suggests that the neuronal receptor GPC5 signals through may determine this specificity. Neuronal receptors that are located at synapses have been identified for GPC family members produced by both astrocytes and neurons. These include presynaptic PTPRD and PTPRS [9, 14, 42], postsynaptic LRRTM4 [12, 13], and postsynaptic GPR158 [42]. In the hippocampus neuronal GPC4 binds postsynaptic GPR158 to induce presynaptic differentiation, and loss of GPR158 leads to immature synaptic morphology specifically at mossy fiber synapses on CA3 neurons, similar to the thalamocortical phenotype in Gpc5 cKO mice [42]. Future studies should investigate if GPR158 or other candidate receptors are expressed by L4 neurons and responsible for mediating the effects of GPC5.

GPC5 can exist in a membrane attached form via its GPI-anchor, or be cleaved from the membrane and act in a soluble form in the extracellular space [11]. Previous studies of astrocyte GPC4 and GPC6 showed that they were functional in their cleaved form [8], and in cell culture soluble GPC5 protein is able to induce presynaptic specializations showing functionality. Determining the form of astrocytic GPC5 that is active in the brain will give insight into its mechanism of action. For example, both soluble and membrane-bound GPC4 regulate synaptogenesis through PTPRS, whereas only membrane-bound GPC4 can signal through GPR158 [9, 14, 42]. This demonstrates that GPCs can have different mechanisms of action and receptor specificity depending on their form. This may explain why the remaining GPC5 expressed by OPCs is unable to compensate for loss of GPC5 in astrocytes, if for example membrane-bound GPC5 is the dominant form in vivo and requires close association between astrocyte processes and the synapse.

This study identifies astrocyte GPC5 as playing an important role in synapse maturation. We show that GPC5 is necessary for refinement and strengthening of thalamocortical synapses, which has implications for the fidelity of thalamic input to the VC and downstream intracortical circuit maturation. In humans GPC5 has been linked to multiple neurological disorders including schizophrenia, Sanfillipo syndrome type B and Alzheimer’s disease [23–28]. Our findings demonstrate that absence of GPC5 may destabilize axonal terminals making them vulnerable to elimination and synapse loss, which could give insight into their role in neurological disorders, for example Alzheimer’s disease, where GPC5 levels are decreased.

## Acknowledgments

We thank Cari Dowling and Joseph Hash for technical assistance, and Alison Caldwell for initial experiments on GPC5. This work was supported by NIH-NINDS R01 NS089791 to N.J.A., as well as the Pew Foundation and CZI Neurodegeneration Network. This work was supported by the Waitt Advanced Biophotonics Core Facility of the Salk Institute with funding from NIH-NCI CCSG: P30 014195 and the Waitt Foundation. Electron microscopy image processing was supported in part by the grants: NN1 NSF 1707356 and NN2 NSF 2014862. The authors acknowledge the Texas Advanced Computing Center (TACC) at The University of Texas at Austin for providing HPC and visualization resources that have contributed to the research results reported within this paper. This work was supported by the GT3 Core Facility of the Salk Institute with funding from NIH-NCI CCSG: P30 014195, an NINDS R24 Core Grant and funding from NEI. U.M. is supported by NSF NeuroNex Award (2014862) and the Chan-Zuckerberg Initiative Imaging Scientist Award. M.C. is supported by a NASEM Ford Foundation Predoctoral Fellowship. L.S. is supported by NIH-NEI F32 EY033629. I.H.S. is supported by fellowships from the Bright Focus Foundation and Alzheimer’s Association.

## Author contributions

A.P.B., M.C., S.W.N., L.S., I.H.S. and N.J.A. performed experiments and analyzed data. U.M. and

N.J.A supervised experiments. A.P.B. and N.J.A. designed the experiments and wrote the paper, with input from all authors. N.J.A. conceived the project.

## Supplemental Figure Legends

**Figure S1 (related to Figure 1).**
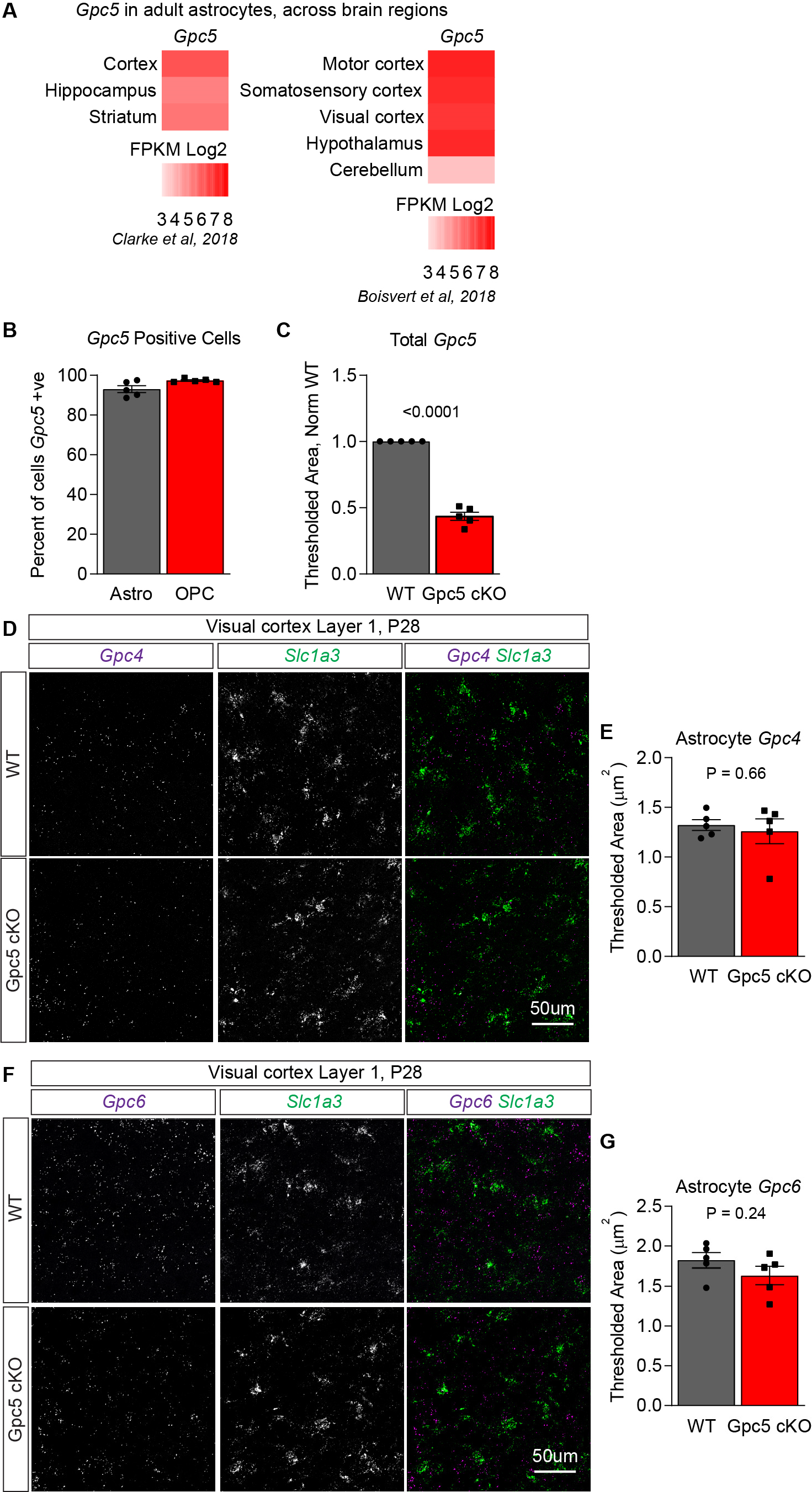
*Gpc5* is expressed throughout the brain by both astrocytes and OPCs. **A.** Expression of *Gpc5* by adult astrocytes across brain regions shows *Gpc5* is enriched in forebrain astrocytes. Data from Clarke et al. 2018 and Boisvert et al. 2018. **B.** *Gpc5* mRNA is expressed by the majority of astrocytes and OPCs within WT VC at P28. Quantification of Figure 1G, colocalization of *Gpc5* with the astrocyte marker *Slc1a3* or the OPC marker *Cspg4*. N=5 mice. **C.** Total *Gpc5* mRNA is decreased by ∼60% in Gpc5 cKO mice. N=5 mice/condition. Graphs mean ± SEM, individual data points mice. Statistics by one sample T-test, P-value on graph. **D-G.** *Gpc4* and *Gpc6* astrocyte mRNA level is unchanged in Gpc5 cKO mice. **D.** Representative images of *Gpc4* mRNA in WT and Gpc5 cKO P28 VC, colocalized with the astrocyte marker *Slc1a3*. **F.** Representative images of *Gpc6* mRNA in WT and Gpc5 cKO P28 VC, colocalized with the astrocyte marker *Slc1a3*. **E,G.** Quantification of D and F respectively. N=5 mice/condition. Graphs mean ± SEM, individual data points mice. Statistics by T-test, P-value on graph.

**Figure S2 (related to Figure 2).**
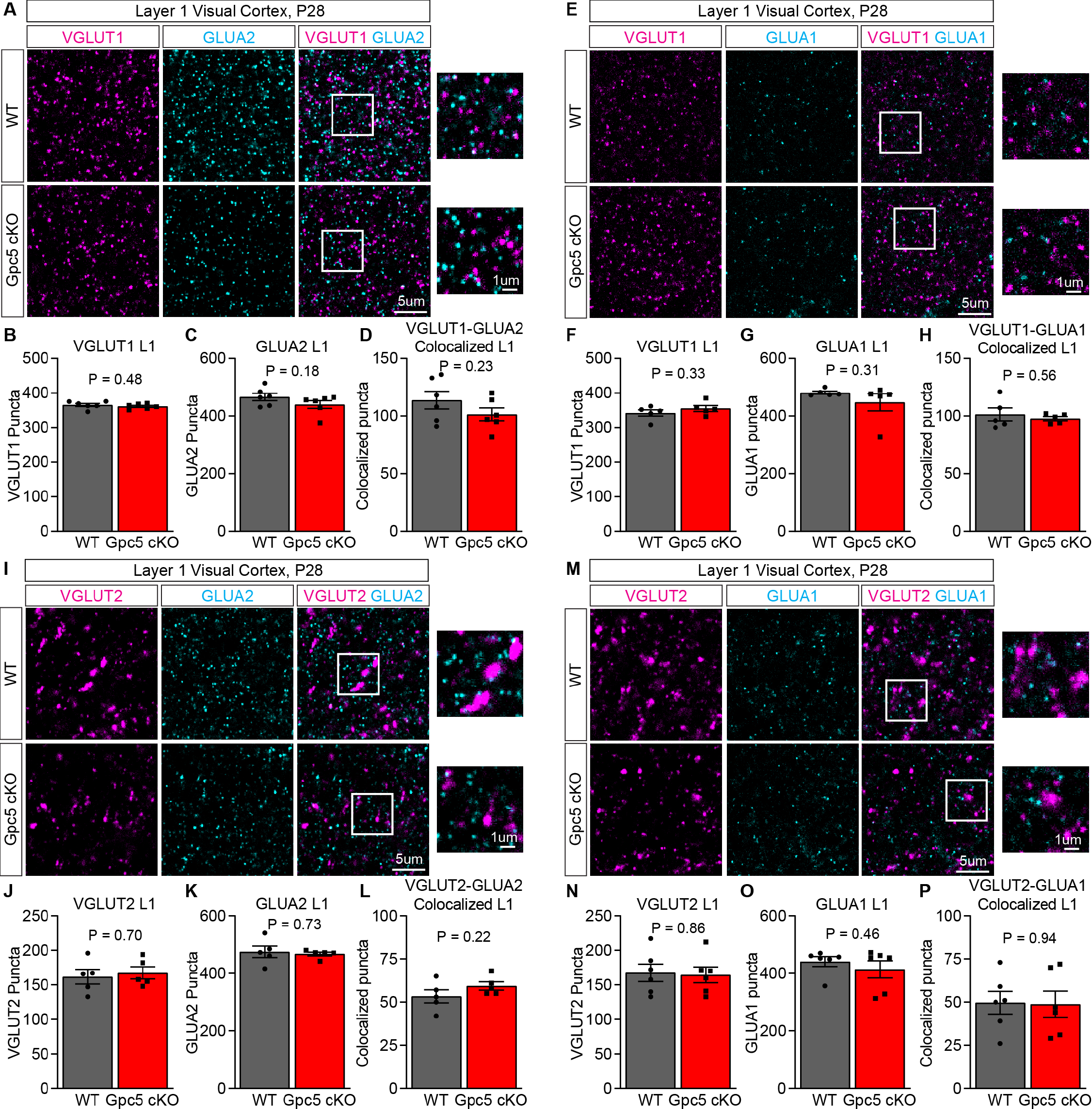
Synapses in Gpc5 cKO mice are immature during the critical period. **A-H.** Intracortical L1 synapses in Gpc5 cKO mice have unaltered AMPAR levels at P28. A. Representative images of immunostaining for intracortical presynaptic marker VGLUT1 and postsynaptic GLUA2 in L1. **B-D.** Quantification of immunostaining, number of VGLUT1 (**B**), GLUA2 (**C**) and colocalized (**D**) puncta shows no change. N=6 mice/condition. **E.** Representative images of immunostaining for intracortical presynaptic marker VGLUT1 and postsynaptic GLUA1 in L1. **F-H.** Quantification of immunostaining, number of VGLUT1 (**F**), GLUA1 (**G**) and colocalized (**H**) puncta shows no change. N=5 mice/condition. **I-P.** Thalamocortical synapses in L1 have unaltered AMPAR level in Gpc5 cKO mice at P28. I. Representative images of immunostaining for thalamocortical presynaptic marker VGLUT2 and postsynaptic GLUA2 in L1. **J-L.** Quantification of immunostaining, number of VGLUT2 (**J**), GLUA2 (**K**) and colocalized (**L**) puncta shows no change. N=5 mice/condition. **M.** Representative images of immunostaining for thalamocortical presynaptic marker VGLUT2 and postsynaptic GLUA1 in L1. **N-P.** Quantification of immunostaining, number of VGLUT2 (**N**), GLUA1 (**O**) and colocalized (**P**) puncta shows no change. N=6 mice/condition. Graphs show mean ± SEM, individual data points represent mice. Statistics by 2-sided T-test, P-value on graph.

**Figure S3 (related to Figure 3.**
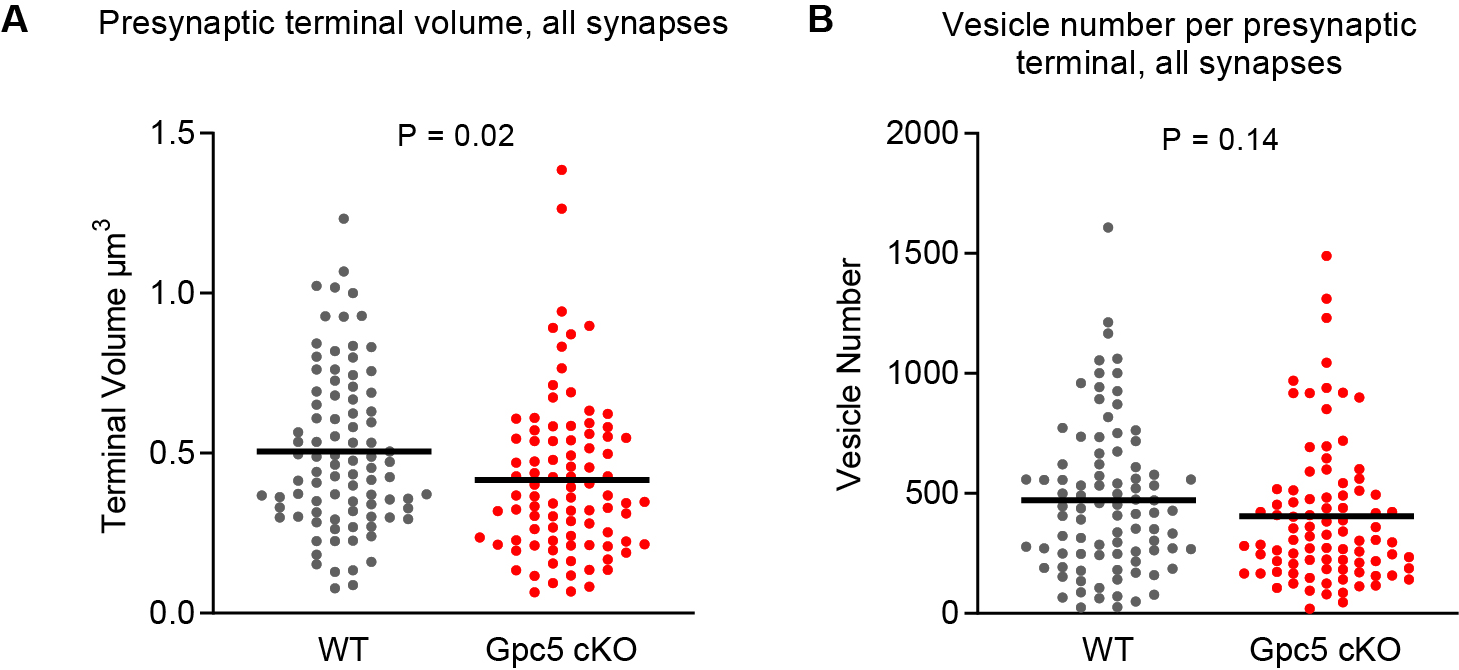
Thalamocortical synapses are structurally immature in Gpc5 cKO mice. **A.** Volume of all presynaptic thalamocortical axonal boutons, combining monosynaptic and multisynaptic boutons, are decreased in Gpc5 cKO mice compared to WT. **B.** The number of synaptic vesicles in all thalamocortical axonal boutons, combining monosynaptic and multisynaptic boutons, is not altered in Gpc5 cKO mice compared to WT. A,B graphs show mean, individual data points represent presynaptic boutons. N=90 presynaptic boutons per condition, statistics by T-test, P-values on graph.

**Figure S4 (related to Figure 4).**
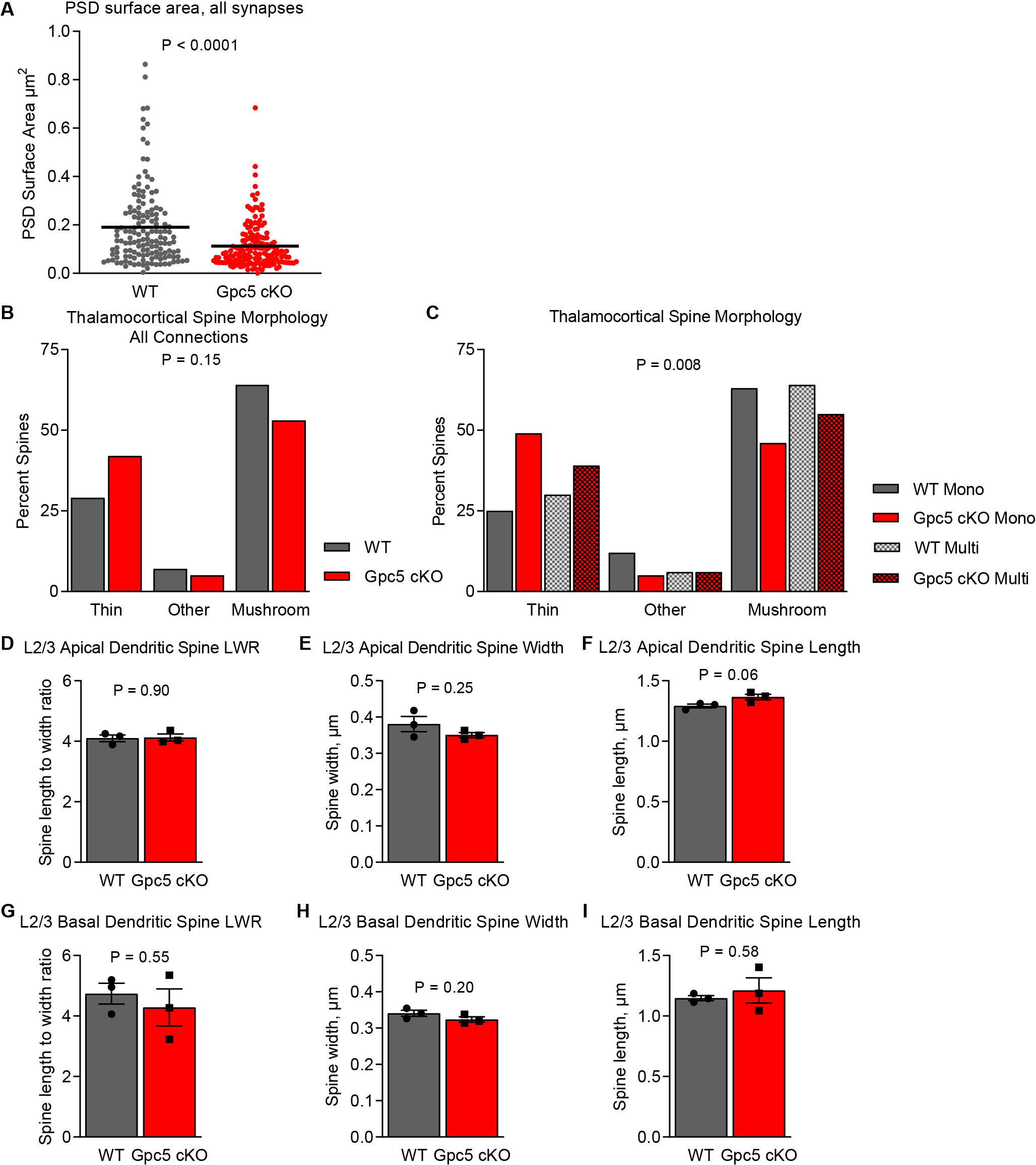
Gpc5 cKO mice show altered postsynaptic structure at thalamocortical synapses. **A.** Surface area of PSD at thalamocortical synapses in Gpc5 cKO mice is decreased when analyzing monosynaptic and multisynaptic connections combined. Graphs show mean, individual data points represent PSDs. N=90 presynaptic boutons and associated PSDs per condition, statistics by T-test, P-value on graph. **B,C.** Dendritic spine structure at thalamocortical synapses is shifted towards a more immature state in Gpc5 cKO mice. Prevalence of thin spines is increased and mushroom spines decreased when analyzing monosynaptic and multisynaptic connections separately (**C**) but not when combined (**B**). Other category includes stubby spines and synapses directly onto the dendritic shaft. Data presented as percentage of spines in each category, spines identified as opposed to presynaptic boutons analyzed in Figure 3. N=90 presynaptic boutons and associated spines per condition, statistics by Chi-square test, P-value on graph. **D-I.** Spines on L2/3 neurons are unaltered in Gpc5 cKO mice. Quantification of Figure 4F. No change in the length (**F,I**), width (**E,H**) or length to width ratio (LWR) (**D,G**) of dendritic spines on secondary apical and basal dendrites in Gpc5 cKO. N=3 mice/condition. Graphs mean ± SEM, individual data points mice. Statistics by T-test, P-value on graph.

**Figure S5 (related to Figure 5).**
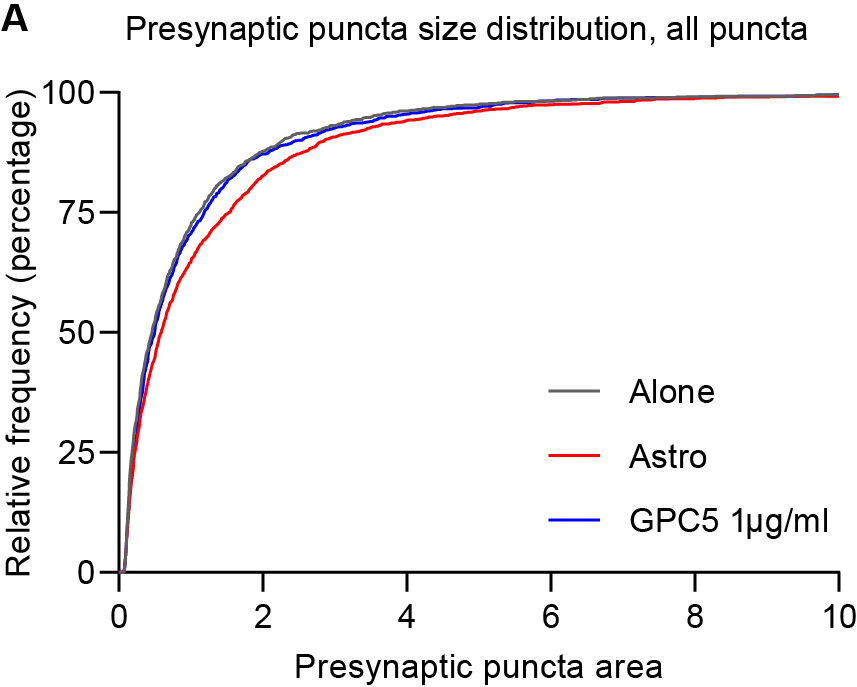
GPC5 is sufficient to induce presynaptic specializations. Treating RGC neurons in culture with soluble GPC5 is sufficient to increase the number of presynaptic terminals without increasing terminal size. **A.** Presynaptic terminal size distribution, all puncta, taken from N=110 cells per condition from 4 experiments.

## METHODS

### Animals

All animal experiments were approved by the Salk Institute IACUC.

***Rats:*** Sprague Dawley rats (Charles River stock number 001) were housed with a 12-hour light/dark cycle in the Salk Institute animal facilities. Rats were provided access to food and water ad libitum. For astrocyte and neuron cell culture experiments, both sexes were used.

***Mice:*** Mice were housed with a 12-hour light/dark cycle in the Salk Institute animal facilities. Mice were provided access to food and water ad libitum. Mice of both sexes were used.

*Astrocyte-specific Glypican 5 conditional knock out mice:* To selectively remove Gpc5 from astrocytes, Gpc5 floxed mice were crossed to B6.Cg-Tg(Gfap-cre)73.12Mvs/J (Jax stock number 012886). Gpc5 floxed mice were generated by KOMP/MMRRC/EUCOMM as conditional ready mice. Gpc5 strain was received as the tm1a allele (C57BL/6N-Atm1Brd Gpc5tm1a(KOMP)Wtsi/, MMRRC Stock #: 047921-UCD) and crossed with mice expressing Flp recombinase (B6.129S4-Gt(ROSA)26Sortm1(FLP1)Dym/RainJ, Jax stock number 009086) to generate Gpc5tm1c(KOMP)Wtsi (UC Davis KOMP repository, project ID CSD76974). All experiments were performed using Gpc5 flox/flox cre negative (WT) and cre positive (cKO) littermate pairs.

### Tissue collection and preparation

Mice were anesthetized with an intraperitoneal injection of 100mg/kg Ketamine (Victor Medical Company) and 20 mg/kg Xylazine (Anased) prior to intracardial perfusion. For collection of fresh frozen tissue used for single molecule fluorescent in situ hybridization (smFISH) experiments, mice were transcardially perfused with 10 mls PBS. Collected brains were embedded in OCT (Scigen 4583), frozen in dry ice/ethanol, and stored at -80°C. For collection of fixed brains used for immunohistochemistry experiments, mice were transcardially perfused with 10 mls PBS followed by 10 mls 4% PFA. Collected brains were placed in 4% PFA overnight at 4°C, washed 3 times in PBS, and cryoprotected in 30% sucrose at 4°C before being embedded in TFM (General data healthcare TFM-5), frozen in dry ice/ethanol and stored at -80°C. For cell fill experiments, mice were transcardially perfused with oxygenated aCSF (in mM: NaCl 126, NaHCO_3_ 26, Glucose 10, KCl 2.5, MgCl_2_ 2, NaH_2_PO_4_ 1.25, CaCl_2_ 2, pH 7.4) at 34°C for 30 seconds followed by 34°C 4% PFA for 15 minutes. Brains were collected and immediately sectioned on a vibratome. For electron microscopy (EM) experiments, mice were transcardially perfused with oxygenated aCSF (in mM: NaCl 126, NaHCO_3_ 26, Glucose 10, KCl 2.5, MgCl_2_ 2, NaH_2_PO_4_ 1.25, CaCl_2_ 2, pH 7.4) with 20U/mL Heparin (Sigma Aldrich H3393) at 34°C for 30 seconds followed by 75 mls of 0.15 M Cacodylate buffer, 2.5% Glutaraldehyde, 2% PFA, 4mM CaCl_2_ warmed to 37°C. Brains were collected and stored overnight at 4°C in 0.15 M Cacodylate buffer, 2.5% Glutaraldehyde, 2% PFA, 4mM CaCl_2_. Prior to sectioning on a vibratome, brains were washed three times in ice cold 0.15M Cacodylate buffer with 4mM CaCl_2_.

### Single molecule fluorescent in situ hybridization (smFISH)

Gpc5 WT and cKO littermate mice were used at P28 to analyze *Gpc5* cell-type expression and efficiency of GPC5 removal from astrocytes, and at P28 and P120 to analyze *Arc* expression in the BZ of VC following monocular enucleation. Fresh frozen, 18 µm coronal sections (3.4 mm posterior to Bregma) were obtained using a cryostat (Hacker Industries OTF5000), or sagittal sections for Figure 1D. smFISH RNAscope (ACDbio 320850) was performed following manufacturer’s instructions for fresh frozen tissue. Slides were frozen at -20C for 20 minutes, followed by 15 minutes in 4% PFA at 4C. Sections then underwent dehydration via 5 minute washes in 50%, 75%, and 100% (x2) ethanol. Following dehydration, sections were incubated with Protease 3 (P28) or Protease 4 (P120) for 15 minutes at room temperature and then washed 2 times in PBS. Slides were incubated with target probes for 2 hours at 40C followed by 3 amplification steps and 1 detection step with RNAscope wash buffer rinses between each step. Sections were mounted with SlowFade gold antifade with DAPI (Thermo Fisher S36939) and coverslip applied (22 mm × 50 mm, 1.5 thickness) and sealed with clear nail polish. Slides were imaged within 1 day or stored at -20C.

Probes used were GPC4 (ACDbio 442821), GPC5 (ACDbio 442831), GPC6 (ACDbio 453301), SLC1A3/GLAST (ACDBio 430781-C2), CSPG4 (ACDbio 404131-C3), and ARC (ACDbio 316911). A negative 3-plex probe (ACDbio 320871) was used as a negative control to determine the level of background signal. Probes for GPC4, GPC5, GPC6 and ARC were imaged in channel 550; SLC1A3/GLAST was imaged in channel 488; CSPG4 was imaged in channel 647.

To determine GPC expression layers 1 to 6 of the VC were imaged using a 20X objective on a Zeiss LSM710 confocal microscope at 2048×2048 pixels as 2µm z-stacks (3 slices). Representative images are maximum intensity projections of the z-stack. Quantification of smFISH signal was performed using an ImageJ macro [15]. Images were made into maximum intensity projections and astrocytes and OPCs identified by their respective probes, with an region of interest (ROI) drawn around each astrocyte or OPC cell body. The probe of interest (GPC4, GPC5 or GPC6) channel was thresholded in the same way for all images, cell-type ROIs were applied, and the thresholded area of probe signal recorded for each ROI. A minimum of 5 littermate pairs of mice, and 3 sections per mouse were imaged. Data in graphs presented as average per mouse.

For *Arc* experiments entire coronal sections containing the BZ of VC were imaged with a 10x objective, as 16-bit images on a Zeiss Axio Imager.Z2 fluorescent microscope with 10% tile overlap. A minimum of 5 littermate pairs of mice, and 4 sections per mouse were imaged. Data in graphs presented as average per mouse.

### Immunohistochemistry and synaptic puncta analysis

Littermate pairs of Gpc5 WT and cKO mice were used for immunohistochemistry experiments at P28 and P120. Coronal sections (20µm) containing the BZ of VC were cut from PFA fixed mouse brains on a cryostat, mounted on Superfrost Plus micro slides (VWR 48311-703), and immediately processed for immunohistochemistry. Sections were placed in a RT humidified chamber to be blocked and permeabilized for 1 hour in 5% goat serum and 0.3% Triton X-100 in PBS. Sections were incubated with primary antibodies in a humidified chamber overnight at 4°C. Primary antibodies were diluted in 5% goat serum, 0.3% Triton X-100, and 100mM lysine in PBS. Primary antibodies used: rabbit anti-GLUA1 (Millipore AB1504) 1:500; rabbit anti-GLUA2 (Millipore AB1768-I) 1:500; guinea pig anti-VGLUT1 (Millipore AB5905) 1:1000; guinea pig anti-VGLUT2 (Millipore AB2251) 1:1000. Sections were washed 3×5 minutes in PBS, then incubated with secondary antibodies in a humidified chamber at RT for 2 hours. Secondary antibodies were diluted in 5% goat serum, 0.3% Triton X-100, and 100mM lysine in PBS. Secondary antibodies used: goat anti-rabbit Alexa 488 (Thermo Fisher Scientific A11073) 1:500; goat anti-guinea pig Alexa 594 (Thermo Fisher Scientific A11032) 1:500. Sections were incubated with secondary antibodies alone as a negative control. Sections were washed 3×5 minutes with PBS. SlowFade gold antifade mountant with DAPI (Thermo Fisher Scientific S36939) was applied to each section and a coverslip (22 mm × 50 mm 1.5 thickness) placed on top and sealed with clear nail polish.

Images were acquired on a Zeiss LSM-880 confocal microscope using a 63x oil immersion objective (1.4NA) as 16-bit images, 1420×1420 pixels, 0.08µm × 0.08µm pixel size, as a z-stack of 8 slices with a total thickness of 2.68µm. For VGLUT1 and GLUA1/2 co-staining images were taken in VC L1 and L2/3. For VGLUT2 and GLUA1/2 co-staining images were taken in VC L1 and L4. Imaging conditions were determined based on the WT condition and applied to the cKO acquired in the same session.

Synaptic staining images were analyzed using IMARIS software (Bitplane) to determine individual puncta number (VGLUT, GLUA) and synapse number (colocalized VGLUT and GLUA1). Four 25µm × 25µm ROIs were selected from within each image for analysis. Each z-stack was viewed as a 3D image and a Gaussian filter of 0.0725µm applied. Puncta were selected by manually thresholding the image and defined using the spots tool as spheres with a set diameter: GLUA1 0.4µm, GLUA2 0.4µm, VGLUT1 0.4µm, VGLUT2 0.5µm. Synapses were defined as colocalization of presynaptic puncta (VGLUT1 or VGLUT2) and postsynaptic puncta (GLUA1 or GLUA2) using the spots colocalization function, measuring a distance of 0.7µm from center to center of each spot. Volume of VGLUT2 puncta was measured using the surface tool, thresholded to capture all puncta defined by the spots tool. All analysis was done blind to genotype. A minimum of 5 littermate pairs and 3 sections per animal were imaged and analyzed. Data in graphs presented as average per mouse. Example images are from a single confocal plane.

### Cell fills

Littermate pairs of Gpc5 WT and cKO mice were used for experiments at P28. Coronal sections (200µm) of lightly PFA fixed tissue were cut on a vibratome in ice cold PBS and stored in 4°C PBS for up to 48 hours. Slices were placed in RT PBS on the stage of a Zeiss microscope, and pyramidal neurons in VC L2/3 identified and soma impaled with a sharp micropipette (100-400 MΩ) backfilled with 10mM Alexa 488 (Thermo Fisher A10436) in 200 mM KCl. Dye was injected by applying a 1.5 V negative pulse for 5-10 minutes until the cell was filled. After filling, slices were placed in 4°C 4% PFA for 30 minutes. Slices were mounted on slides with SlowFade gold antifade mountant with DAPI (Thermo Fisher Scientific S36939), coverslip applied (22 mm × 50 mm 1.5 thickness) and sealed with clear nail polish. Slides were prepared for slices by applying a thick clear nail polish boundary to prevent coverslips from crushing slices.

Images were acquired on a Zeiss LSM-880 confocal microscope using a 63x oil immersion objective as 16-bit, 1548 × 1548 pixel area, 0.08µm × 0.08µm pixel size images. Exposure parameters were established based on WT samples and all sections were imaged in a single session. A z-stack (0.19µm step size) spanning the entire dendrite was taken for each cell, and both basal and apical secondary dendrites were imaged. Spine analysis was performed using NeuronStudio software [43]. A 15µm segment of secondary apical or basal dendrite was selected, at the first branch point, and the number of spines, spine neck length and spine head diameter were measured and classified according to [44]. A minimum of 5 littermate pairs of mice and 3 cells per mouse were imaged. Data in graphs presented as average per mouse.

### APEX2 injection and electron microscopy

To label thalamocortical projections for EM reconstruction, AAV9-COX4-dAPEX2 was injected into the dLGN of WT and cKO mice at P14 (2 littermate pairs). pAAV-COX4-dAPEX2 was a gift from David Ginty (Addgene plasmid #117176; http://n2t.net/addgene:117176; RRID:Addgene_117176) [34]. Packaging in AAV9 was performed by the Salk Viral Vector core facility (GT3) at a concentration of 2×10^14^ vg/mL. Mice were anesthetized with oxygenated isoflurane (2-3%) and injection was done with a Nanoject pressure injection system. Virus was diluted to 3×10^12^ vg/mL and injected at coordinates 2.0 mm posterior from bregma, 1.9 mm lateral from the midline, and 2.9 mm below the pia, with a total of 150nL of virus delivered at a rate of 2nL per second. Following 2 weeks of expression, mice were collected at P28 as described above and the brain was processed for electron microscopy as described with some modifications [34, 45]. Materials used for processing samples for EM were sourced from Electron Microscopy Sciences unless otherwise indicated. All steps were performed at ice cold temperatures unless otherwise indicated.

Brains were mounted on a Leica VT1000S vibrating microtome in cacodylate buffer, and 100µm coronal sections containing the primary VC collected in 6 well plates and washed 2×10 minutes in cacodylate buffer supplemented with 50mM glycine, followed by 1×10 minutes in cacodylate buffer. A 10X diaminobenzidine (DAB) tetrahydrochloride solution was freshly prepared by dissolving 50mg of DAB in 0.1 M HCl at room temperature prior to tissue processing. Sections were then incubated in DAB solution (final concentration of 0.3 mg/mL DAB in cacodylate buffer) for 30 minutes in the dark. After 30 minutes, 10µL/mL of cacodylate supplemented with 0.3% H_2_O_2_ was added directly to the DAB solution (final H_2_O_2_ concentration of 0.003%) and swirled extensively to initiate the peroxidase reaction which was allowed to proceed for 1 hour in the dark. Slices were evaluated for reaction product and washed 3×10 minutes in cacodylate buffer and then further post-fixed overnight in cacodylate buffer with 3% glutaraldehyde.

The following day sections were rinsed 2×10 minutes in cacodylate buffer with 50mM glycine followed by 1×10 minutes in cacodylate buffer and transferred to a petri dish filled with ice cold cacodylate buffer for photography and microdissection. 2mm wide strips spanning from the cortical surface to the corpus collosum were collected into scintillation vials for further processing. Samples were stained with reduced osmium (1% osmium tetroxide and 1.5% potassium ferrocyanide in cacodylate buffer) for 1h at room temperature, then rinsed 5x3 minutes with ice cold water and left in 1% aqueous uranyl acetate at 4°C overnight. Samples were then serially dehydrated in ice cold aqueous ethanol solutions of ascending concentrations, before 3×10 minute incubations with absolute ethanol at room temperature. Samples were then infiltrated with ascending concentrations of Durcupan resin in absolute ethanol at room temperature (3:1, 4h; 1:1, 4h; 1:3, overnight) before 2x4h incubations rotating in pure resin. Samples were embedded with fresh resin and paper labels in silicon molds, with the tissue oriented *en face* to the block face and polymerized for 60h at 65°C in an oven.

Serial sections were collected onto silicon wafers as described with some modifications [46]. Briefly, the block was trimmed using a 90° diamond trimming knife (Diatome) on an ultramicrotome (Leica UC7) to a trapezoidal frustum of roughly 150×400µm which included the region from the cortical surface to deep cortical layers. A silicon chip (35x7mm; University Wafer, Boston, MA) was hydrophilized in a plasma cleaner (Harrick) immediately preceding partial immersion in the water boat of a Histo knife (Diatome) mounted on the ultramicrotome. Ribbons of 150-200 serial sections of thicknesses of approximately 55nm were cut with 4 drops of pure ethanol in the water boat and an ionizing instrument (Leica EM Crion) activated and oriented towards the cutting edge of the knife from above. When ribbons of sufficient quality and length were generated, they were released from the knife edge using a single-eyelash brush and carefully positioned over the chip. The water level was then slowly lowered using a peristaltic pump, and sections were allowed to dry down on the silicon substrate. Chips were briefly dried on a slide on a hot plate set to 60°C.

Chips were mounted on aluminum stubs using carbon sticky tabs and loaded into a scanning electron microscope (SEM; Zeiss Sigma VP) equipped with a sensitive backscatter detector (Gatan), as well as extended raster scanning capabilities and a control system designed for serial section imaging workflows (ATLAS5, FIBICS). Low resolution image maps of the ribbon of serial sections were collected, and a mid-resolution map of a central section was generated for reference. From this image, a region of interest (ROI) from VC L4 of 50×50µm was selected from between 250-350µm from the cortical surface that [1] had DAB+ terminals; [2] was not obstructed by blood vessels or somata; [3] was free from obvious debris throughout the series as assessed from the low-resolution map. This region was identified at one end of the ribbon of sections, and the ROI was imaged at high resolution (pixel size: 2nm; dwell time: 6µs; EHT: 3kV; aperture: 30µm; working distance: 8-9mm) on every section in the ribbon.

Image stacks were collated and rigidly aligned using TrakEM2 in Fiji and cropped to a minimum continuous cube of roughly aligned data with minimal padding [47]. Fine stack alignment was accomplished using SWiFT-IR as deployed on 3DEM.org using the TACC compute resource Stampede 2 [48, 49]. The well aligned data was then imported into VAST Lite (VAST) for annotation and analysis [50]. Briefly, axons with DAB+ mitochondria and their corresponding postsynaptic partners were identified and manually segmented in VAST. Volume of terminals, vesicle cloud size, and PSD surface area were determined by individually segmenting structures and using VASTTOOLS MATLAB toolkit. To categorize post synaptic targets, all post synaptic structures synapsing with a target axonal terminal were segmented. 3D reconstructions of the segmented post synaptic structures were then determined to be either mushroom, thin, stubby, or shaft. Spines were categorized based on visual inspection of morphology. Mushroom spines were identified by the presence of a defined head and neck; thin spines were categorized as long filopodia like structures with no defined head; stubby spines were identified as short structures with no definable neck; shaft synapses occurred directly on the dendritic shaft. Data in graphs presented per presynaptic bouton or per postsynaptic spine. Visualizations of reconstructions were produced using the Neuromorph add-on in Blender 2.7 (Blender Foundation; blender.org) [51].

### Monocular enucleation and *Arc* analysis

Littermate pairs of Gpc5 WT and cKO mice were used for experiments at P28 (critical period) and P120 (adult). Mice were anesthetized with 2% isoflurane in oxygen and the right eye was removed via transection of the optic nerve. The empty ocular cavity was filled with Gelfoam (Pfizer 031508) and eyelid was sutured closed with 6-0 silk sutures (Henry Schein 101-2636). Erthromycin 0.5% and lidocaine 2% were applied to sutured eyelid. Overnight deprivation (control) mice were collected 12 hours later. 5-day monocularly deprived mice were collected after 5 days. Mice were maintained in a 12-hour light/12-hour dark cycle and collected (as described above) at the end of a 12-hour dark cycle. Mice were exposed to 30 minutes of bright light to induce *Arc* expression in neurons in the VC stimulated by the open eye before tissue collection. Coronal sections (18µm) were made from fresh frozen tissue on a cryostat and smFISH for *Arc* and imaging performed as described above. The width of the activated binocular zone was measured by analyzing the width of the *Arc* signal in VC L2/3 contralateral to the deprived eye performed using the measure tool in Zen blue edition software (Zeiss). A minimum of 5 littermate pairs and 4 sections per mouse were analyzed. Data in graphs presented as average per mouse.

### Cell Culture

#### Retinal ganglion cell neuron culture

Retinal ganglion cell neurons (RGCs) were isolated from P5-P7 Sprague Dawley rat retinas using immunopanning as described [9, 52]. Cells were plated at a density of 30,000 cells/well on glass coverslips (12mm diameter, Carolina Biological Supply 633029) treated with poly-D-lysine (Sigma P6407) and laminin (R&D 340001001) and grown in 24-well plates. RGCs were cultured in growth media containing: 50% DMEM (Thermo Fisher 11960044), 50% Neurobasal (Thermo Fisher 21103049), Penicillin-Streptomycin (Thermo Fisher 10437028), Glutamax (Thermo Fisher 35050-061), sodium pyruvate (Thermo Fisher 11360-070), N-acetyl-L-cysteine (Sigma A8199), insulin (Sigma I1882), triiodo-thyronine (Sigma T6397), SATO (containing: transferrin (Sigma T-1147), BSA (Sigma A-4161), progesterone (Sigma P6149), putrescine (Sigma P5780), sodium selenite (Sigma S9133)), B27 (Thermo Fisher 17504044), BDNF (Peprotech 450-02), CNTF (Peprotech 450-13), and forskolin (Sigma F6886). RGCs were cultured in a humidified incubator at 37°C and 10% CO_2_ with a half-media change every 3 days.

#### Cortical astrocyte culture

Cortical astrocytes were isolated and cultured as described from P1-P2 Sprague Dawley rats [15, 53]. After isolation astrocytes were plated in 15cm cell culture plates coated with poly-D-lysine (Sigma P6407) at 3 million cells/plate and maintained at 37°C and 10% CO2. Astrocyte culture medium was DMEM (Thermo Fisher 11960044) with 10% heat-inactivated fetal bovine serum (Thermo Fisher 10437028), Penicillin-Streptomycin (Thermo Fisher 10437028), Glutamax (Thermo Fisher 35050-061), insulin (Sigma I1882), sodium pyruvate (Thermo Fisher 11360-070), hydrocortisone (Sigma H0888), N-acetyl-L-cysteine (Sigma A8199). For RGC feeder layer treatment astrocytes were plated on cell culture inserts for use in a 24-well plate (Thermo Fisher 353104) at 50,000 cells/insert. Before addition to wells containing RGCs, inserts were washed 3 × with 34°C DPBS to remove astrocyte growth medium and switched to RGC growth medium.

#### RGC neuron treatment

RGCs were cultured for 7-10 days in full RGC growth media to allow neurite outgrowth, prior to treatment for 6 days with astrocytes or purified GPC5. There were 3 conditions: RGCs alone (negative control), RGCs with a feeder layer of astrocytes (positive control), RGCs + recombinant mouse GPC5 at 1µg/ml (R&D 2689-G5-050/CF, resuspended in DPBS at 0.1µg/µl).

#### Synaptic staining cultured RGC neurons

RGCs were washed 3 × 5 minutes with 34°C DPBS, fixed in 4% PFA at 34°C for 10 minutes, washed 3 × 5 minutes in PBS, blocked and permeabilized for 30 minutes at RT in 50% antibody buffer (NaCl 150mM, Tris Base 50 mM, BSA 1%, L-Lysine 100 mM), 50% goat serum, and 0.2% Triton X100, washed 1 × 5 minutes with PBS. Primary antibodies were diluted in antibody buffer with 10% goat serum and incubated over night at 4°C: mouse anti-PSD95 (Pierce MA1-045) 1:500, rabbit anti-VGLUT2 (Synaptic Systems 135-403) 1:1000. The next day RGCs were washed 3 × 5 minutes with PBS and incubated with secondary antibodies: goat anti-mouse Alexa 488 (Thermo Fisher A11029) 1:1000, and goat anti-rabbit Alexa 594 (Thermo Fisher A11037) 1:1000 at RT for 1 hour, washed 3 × 5 minutes with PBS and coverslips mounted on microscope slides (Fisherfinest 12-544-2) with SlowFade gold antifade mountant with DAPI (Thermo Fisher S36939) and sealed with clear nail polish.

RGCs were imaged on a Zeiss AxioImager.Z2 microscope with a 63x oil immersion objective. Images were acquired at 14 bit, 1388×1040 image size, pixel size 0.102µm × 0.102µm. RGCs were selected in the DAPI (nucleus) channel, then pre and postsynaptic puncta imaged. Exposure settings for each imaging session were established based on the positive control (+astrocytes) and used for each image. Synaptic analysis was carried out using the ImageJ (NIH) puncta analyzer plug-in as previously described [8, 54]. Briefly, thresholds to select pre and postsynaptic puncta were established using the positive control condition (+astrocytes), then applied to all images to select puncta to be considered for colocalization = synapse. VGLUT2 puncta size analysis was carried out in ImageJ. Images of the VGLUT2 channel were thresholded based on the positive control condition (+astrocytes), then the ‘analyze particles’ function used to select thresholded puncta and measure their area. Two or three coverslips per condition and 10 cells per coverslip were imaged, giving 20-30 cells per group per experiment, and the experiment was repeated on 4 separate cultures. Data are presented as combined cells from all 4 experiments, and within each experiment data are normalized to the RGC alone condition.

### Data presentation and statistical analysis

Data in graphs is presented as mean ± S.E.M. along with individual data points representing mice or cells as indicated in the figure legend. Statistical analysis was performed with Prism and the test used indicated in the figure legend. All tests were 2-tailed, and exact P values are reported on the graph. Data were tested for normality of distribution before statistical testing. For 2 samples an unpaired T-test was used for normally distributed data and a Mann-Whitney test for non-normally distributed data. For more than 2 samples an ANOVA with Tukey’s post-hoc test for multiple comparisons was used for normally distributed data, and a Kruskal-Wallis ANOVA on ranks with Dunn’s post-hoc test for multiple comparisons used for non-normally distributed data. To compare categories a Chi-square test was used.

