## Supplementary figures and images for "Astrocyte glypican 5 regulates synapse maturation and stabilization"

### Supplemental Figures

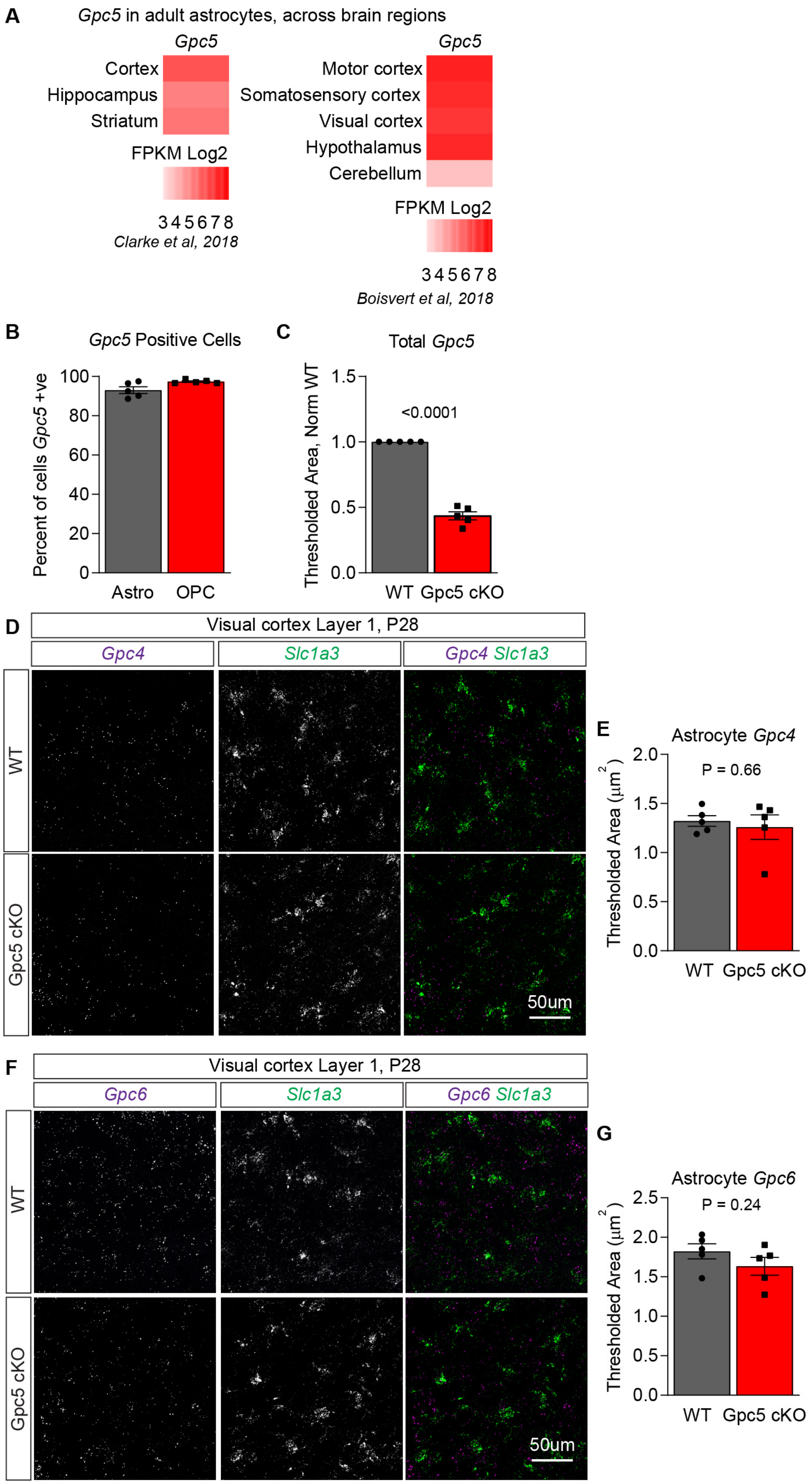

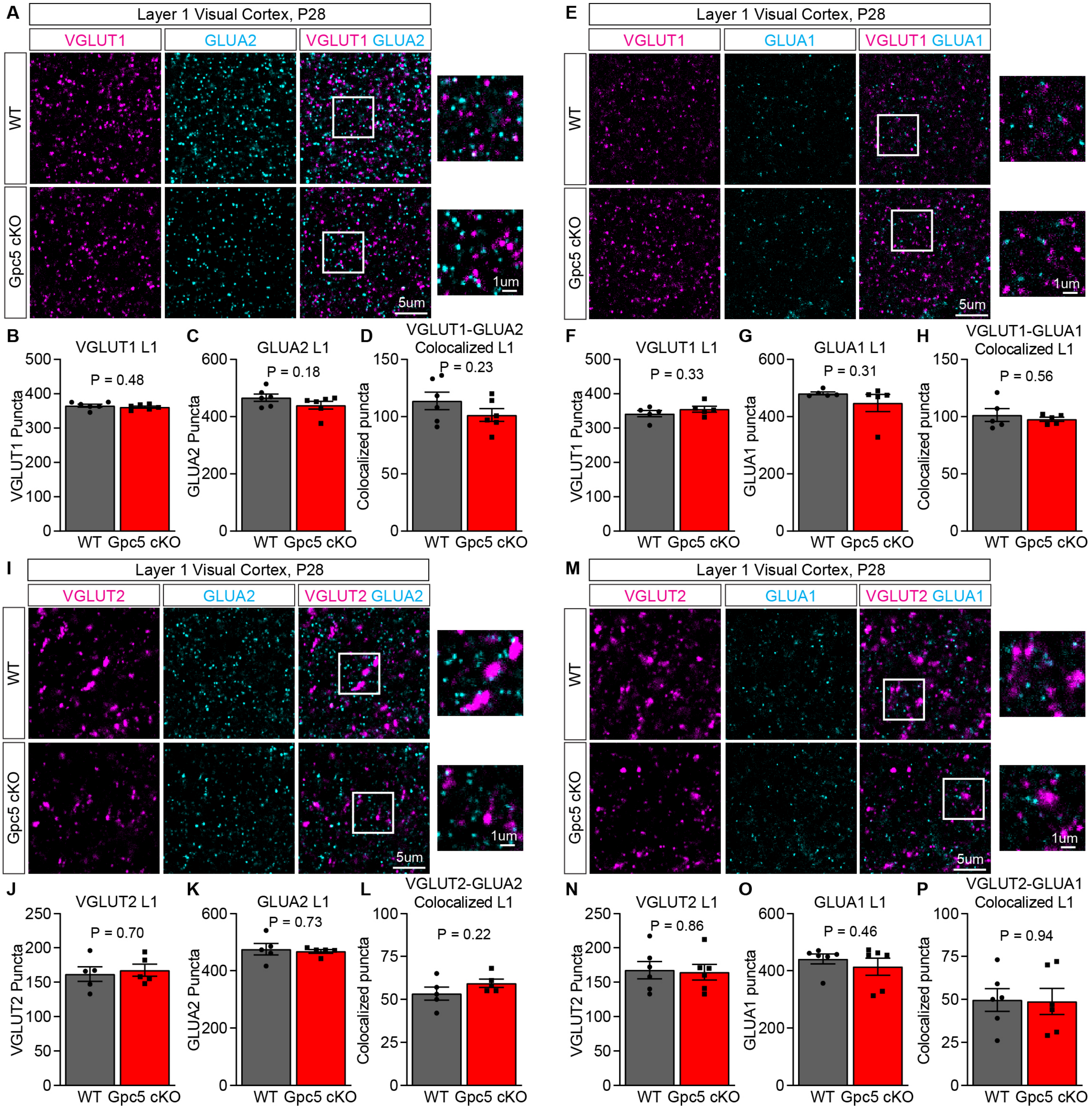

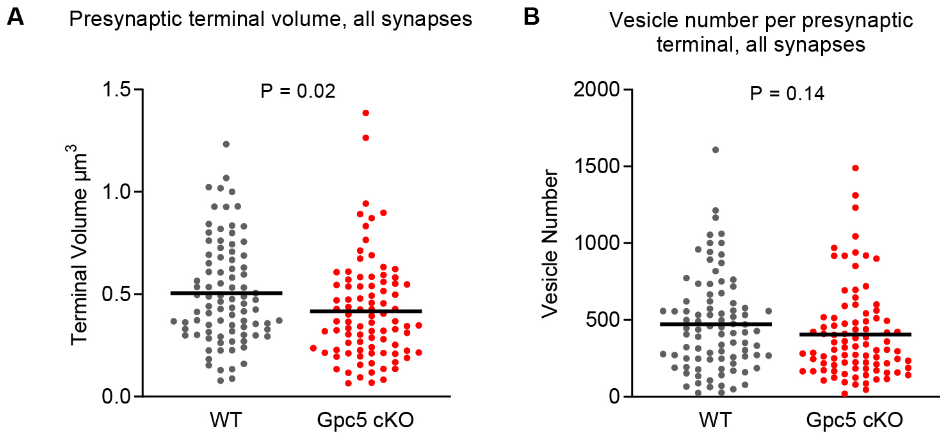

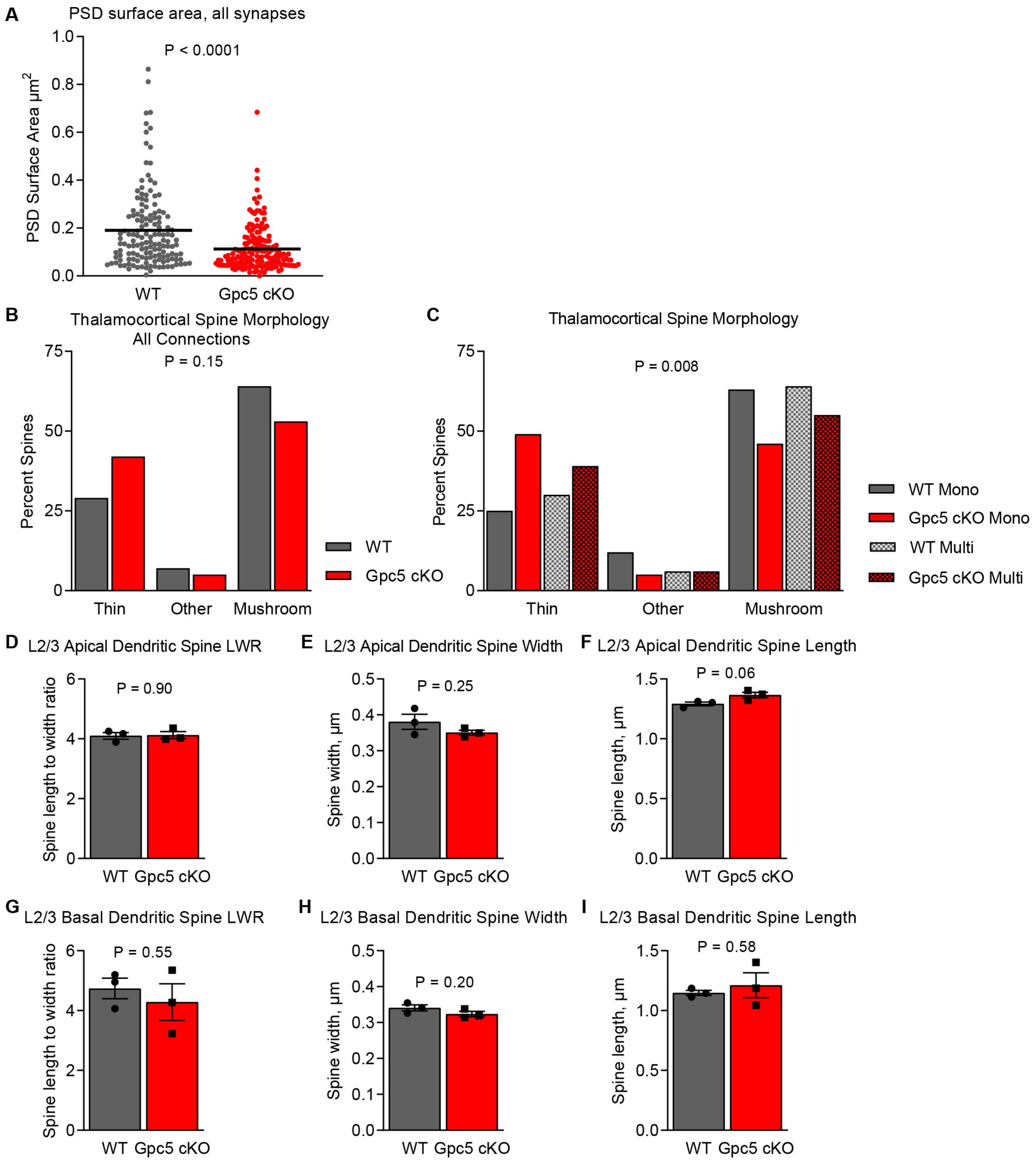

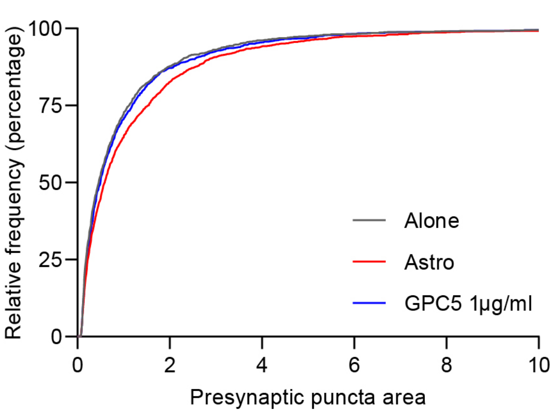
